# Predicting tuberculosis relapse based on 28-day CFU, RS ratio, and/or drug contribution for novel regimens in the relapsing mouse model

**DOI:** 10.64898/2026.07.27.740024

**Authors:** Rob C. van Wijk, Belen P. Solans, Linda Chaba, Sylvie Sordello, Anna M. Upton, Eric L. Nuermberger, Gregory T. Robertson, Nicholas D. Walter, Radojka M. Savic

## Abstract

Treatment shortening in tuberculosis therapy is needed, but testing all novel antibiotic combinations is unfeasible. Especially the tuberculosis relapsing mouse model is time- and resource demanding. Therefore, our objective is to develop a computational model predictive of long-term relapse prevention in mice based on short-term biomarkers, increasing the number of regimens that can be tested and prioritize regimens for further development. The innovative ribosomal RNA synthesis (RS) ratio is utilized to characterize drug effect on Mycobacterium tuberculosis health and activity, together with colony forming units (CFU) in murine lungs. Nine datasets of 58 unique regimens with 843 short-term biomarker and 2,239 long-term relapse observations were leveraged for model development in 3 iterations with external validations. The final model included therapeutic predictors, such as CFU and RS ratio change from baseline, and corrected for experimental conditions, to enable unbiased ranking of regimens between experiments. Model performance was optimal without model structure change despite fully separate model development at each iteration. Final external validation had an area under the receiver operator curve of 0.90. Challenging the model by assessing removal of either biomarker showed that performance of CFU only was similar to CFU and RS ratio once the sterilizing contribution of individual drugs to the regimens was accounted for. New drugs without this contribution quantified could benefit from RS ratio determination to predict relapse. Our predictive model can successfully differentiate between 2-, 3-, and 4-month regimens in the relapsing mouse model based on 4-week data only, supporting acceleration of treatment-shortening regimen development.

**One Sentence Summary:** Our predictive model ranks new drug regimens by tuberculosis relapse prevention based on 28-day CFU and RS ratio, or on CFU only for known drugs.

## INTRODUCTION

Tuberculosis (TB) remains a significant global health challenge, ranking among the top infectious disease killers worldwide (*1*). The persistently high mortality and morbidity associated with TB underscore the urgent need for more effective and shorter-duration treatment regimens. Traditional TB treatments, such as the 6-month regimen for drug-susceptible TB consisting of isoniazid, rifampin, pyrazinamide, and ethambutol (*2*), or the more recent 4-month regimen consisting of isoniazid, rifapentine, pyrazinamide, and moxifloxacin (*3*), are lengthy and often associated with poor patient compliance. Poor adherence increases the risk of treatment failures and the emergence of drug-resistant strains (*4*). Thus, there is a critical demand to innovate and expedite the development of novel, more potent, and shorter TB therapies.

The current TB drug development pipeline is robust, with numerous candidates showing promise in preclinical stages (*5–12*). Combining these numerous candidates into the right regimens with the highest treatment-shortening potential becomes experimentally unfeasible. The translation from preclinical findings to clinical application is already notoriously slow. This delay is primarily due in part to the reliance on long-term endpoints, such as the 9-month relapsing mouse model of TB (RMM). In the RMM, mice are treated with different regimens of interest for different durations, and the number of relapsing mice is observed 12 weeks after the predefined treatment duration concludes (*13–16*). Based on the proportion of mice relapsing after different treatment durations, regimens can be ranked and an indication of treatment shortening compared to the standard of care can be made (*17–19*). Confirmation of improved relapse prevention and treatment shortening potential compared to standard of care in the RMM is considered an important stage-gate in moving a regimen forward to clinical development. But the time-consuming and resource-intensive nature of the RMM impedes the rapid progression and prioritization of potential treatments (*20, 21*). There is a clear need for a more efficient and faster, shorter-term estimation of relapse prevention potential and treatment duration for the larger number of regimens in preclinical development, shifting the RMM from learning to confirming of regimens with the most promising potential (*22*).

Addressing the need for more efficient and faster relapse prevention assessment for novel regimens requires both experimental and computational innovation. Experimentally, there is a need for informative biomarkers that can be quantified in short-term experiments of at most 4 weeks of treatment, which are predictive of relapse prevention potential. The conventional biomarker of *Mycobacterium tuberculosis* colony-forming units (CFU) quantified in lung homogenates during treatment enumerates the burden of bacteria that are capable of multiplying to form a visible colony on a plate. This cumbersome methodology takes an especially long time given the slow replication rate of *M. tuberculosis*, but is nonetheless informative of the ability of the mycobacteria to grow a colony (*23*). However, it is known that CFU alone cannot always distinguish between sterilizing and non-sterilizing regimens (*24*). The presence of one or more sterilizing drugs such as rifamycins, diarylquinolines, or pyrazinamide, in the regimen under assessment can provide additional information on the potential to prevent relapse to distinguish sterilizing from non-sterilizing regimens using quantitative metrics (*25, 26*). For novel drugs, the potential to contribute sterilizing activity to a regimen may not be established through prior relapsing mouse models, and novel biomarkers may provide supporting evidence. One example is biomarkers of pathogen health that measures aspects of mycobacterial physiology (*27, 28*). By quantifying the core physiologic process of rRNA synthesis, the RS ratio provides information complementary to bacterial burden, potentially clarifying why treatments with the same effect on CFU can have different relapse rates when treated for the same duration. Recently, the RS ratio has shown potential as a physiological biomarker to orthogonally inform on relapse prevention (*29–31*). Such a biomarker can provide an early indication of treatment efficacy, thereby accelerating decision-making in the drug development process (*20*). One limitation of the RS ratio which is defined as the ratio of short-lived (ETS1) rRNA by mature (23S) rRNA, is its mechanism-based interaction with antibiotics such as streptomycin or oxazolidinones, which stabilize the short-lived rRNA and therefore impact the ratio by preventing normal ETS1 degradation (*32*).

Innovative computational modelling to develop predictive tools for treatment efficacy has been successful in translational TB drug development, predicting bactericidal activity as well as target site exposure to inform on clinical development (*33–38*). Advanced computational models are necessary to integrate orthogonal biomarkers such as CFU and RS ratio, as well as a quantitative metric of sterilizing contribution of individual drugs to a regimen, into unified predictive frameworks, leveraging diverse mechanistic and physiological signals to improve predictive power of relapse prevention. By combining these complementary data sources, models can estimate relapse probability for regimens not only for tested and observed therapeutic durations but also inter- and extrapolate to previously untested durations, supporting decision-making on optimal treatment duration in drug development. The fundamental statistical nature allows characterization of multiple sources of variability, including experimental conditions, between-experiment variability, and laboratory differences, thereby ensuring unbiased comparison of regimen performance. Moreover, such models facilitate the simulation of novel scenarios to substantially accelerate regimen optimization and translational insight.

In this context, the PAN-TB program emerges as a pivotal initiative, offering a comprehensive data source to refine and validate predictive models. The PAN-TB program integrates data from various preclinical studies and performs comparative experiments, enabling among other objectives the development of robust models that can accurately forecast treatment outcomes based on early biomarkers, regimen composition, and experimental variables (*39–43*). Combining these short-term biomarker data on CFU and RS ratio, with the predictive power of translational modelling, is hypothesized to lead to a translational workflow in which short-term biomarkers obtained on treatment with novel regimens can inform on long-term relapse probabilities for these regimens. If so, the assessment of sterilizing activity could be streamlined into short-term, 4-week treatment experiments in which CFU and RS ratios are determined, which can be done for a larger set of regimens with novel compounds using substantially less mice and less resources, followed by computational prediction of the relapse prevention potential and optimized therapeutic duration as well as a model-informed ranking of regimens. The relapse prevention potential of the highest-ranking regimens could then be confirmed in the formal RMM.

The primary objective of this study is therefore to develop a computational model able to successfully predict the treatment duration necessary to achieve a 95% rate of relapse-free cure (T95) in BALB/c mice, using early-timepoint (≤ 28 days) biomarkers and other experimental variables. Particular focus is on shorter-duration regimens (T95 of less than or equal to 4 months) and the ability of the computational model to distinguish those. The systematic workflow here comprises four key steps: (1) data integration and preparation, (2) model development to establish predictive algorithms including assessment of biomarker contribution and the sterilizing contribution of individual drugs in the regimens, (3) prediction of relapse probability and ranking of regimens based on these models to diagnose and validate them, and (4) external validation based on dedicated validation datasets to ensure the models’ generalizability and accuracy. This framework also includes assessing the capability to forecast the efficacy of novel TB regimens, enhancing the scope of its application (**Figure 1**).

**Figure 1.**
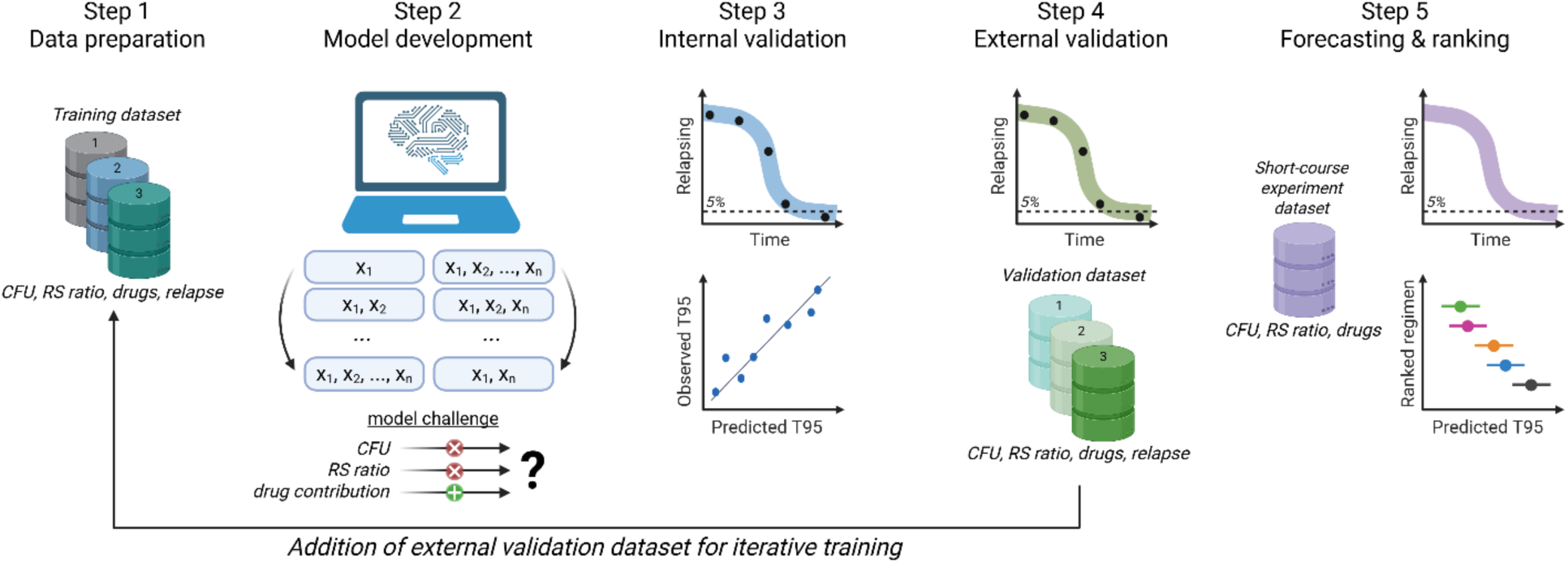
Overview of the analysis methodology. Database of colony forming units (CFU), RS ratio, drugs within the tested regimens, and relapse data for these regimens (step 1) was leveraged to iteratively develop and challenge by sensitivity analysis of a multivariable relapse prediction model (step 2) based on different training (step 3) and external validation (step 4) datasets with increasing size, with diagnostics based on relapse proportion simulations and time until 95% of mice are relapse-free (T95) correlation between prediction and observed. The validated relapse prediction model can be used to forecast relapse probability and T95 for novel regimens based on short-course experiment data on CFU, RS ratio, and/or sterilizing drug contribution (step 5).

By integrating early biomarkers with advanced modeling techniques, our approach aims to significantly shorten the timeline for TB regimen evaluation in mice. This has the potential to transform the TB drug development landscape, enabling faster delivery of new, effective therapies to patients. The findings of this study not only provide a methodological advancement but also have profound implications for future TB treatment strategies, emphasizing the critical role of short-term biomarkers in revolutionizing the development of TB drugs.

## RESULTS

### Large dataset of short-term biomarkers and long-term relapse outcomes

A total of nine BALB/c mouse relapse experiments were included, distributed across three iterations of training and validation datasets as data became available over time (**Table 1, Table S1**). Mice were infected with *M. tuberculosis* strains H37Rv or Erdman and through aerosol or intranasal administration 2 weeks prior to treatment initiation. Colony forming units per lung (CFU/lung) and RS ratio were measured at baseline and at multiple timepoints up to 4 weeks. The long-term outcome metric, relapse-free cure, was defined as 0 CFU/lung 12 weeks after treatment completion. The complete dataset comprised 843 observations for short-term biomarkers of both CFU/lung and RS ratio, and 2,239 relapse outcome observations.

**Table 1.**
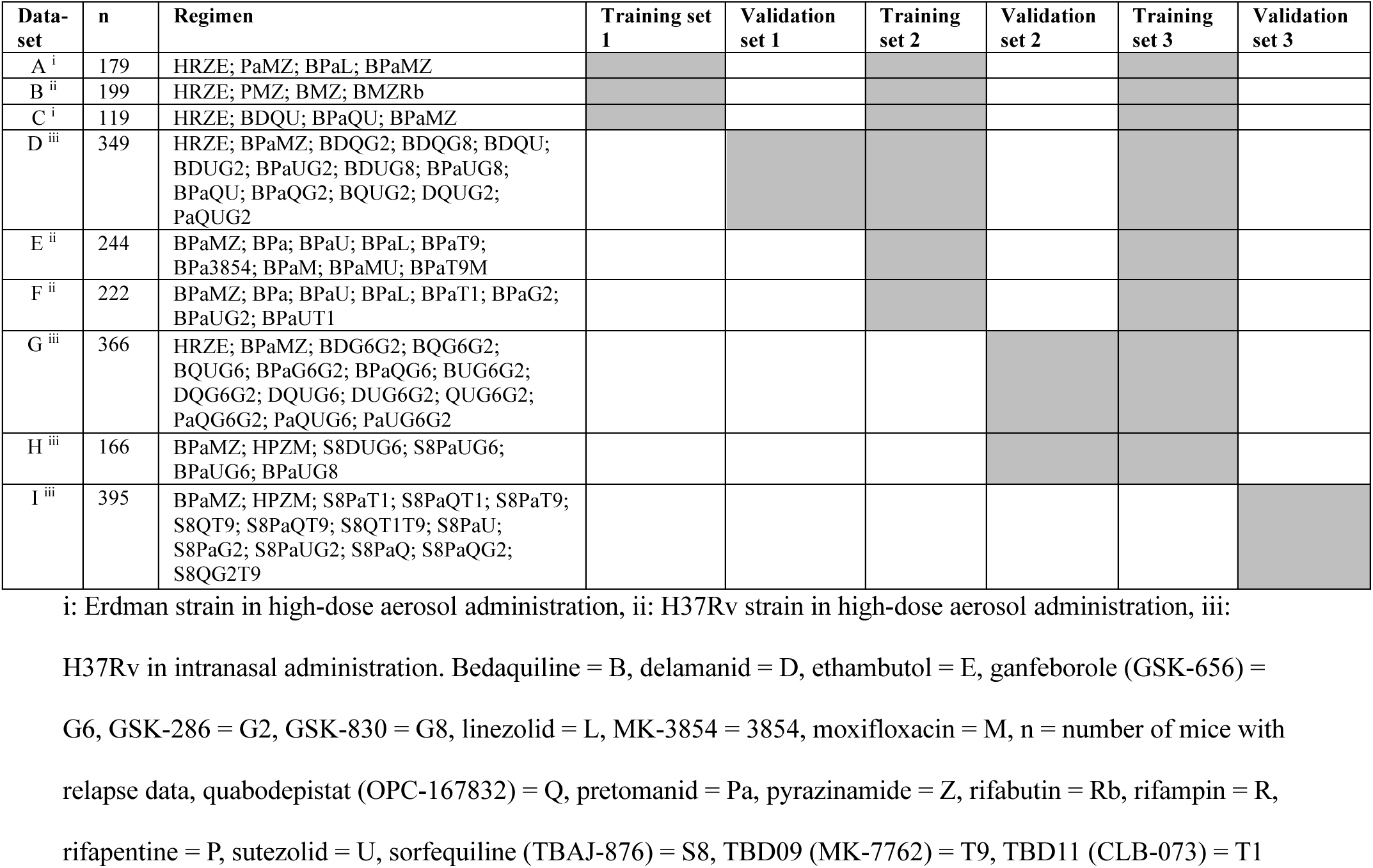
Regimens tested in the BALB/c relapsing mouse model in the included datasets. Dosing in Table S1.

The number of total unique regimens tested was 58, consisting of rifamycins (rifampin, R; rifapentine, P; rifabutin, Rb), diarylquinolines (bedaquiline, B; sorfequiline/TBAJ-876, S8), oxazolidinones (linezolid, L; sutezolid, U; and TBD09/MK-7762, T9), nitroimidazole (delamanid, D; pretomanid, Pa), as well as several novel compounds (the LeuRS inhibitors ganfeborole, G6, and GSK3211830, GSK-830/G8; the DprE1 inhibitor quabodepistat, Q; protein synthesis inhibitor MK-3854, 3854; and the Rv1625c agonists GSK2556286, GSK-286/G2, and TBD11/CLB-073, T1), and ethambutol (E), isoniazid (H), moxifloxacin (M), and pyrazinamide (Z). The presence of oxazolidinones was of particular interest, given their interaction with the innovative RS ratio biomarker. The BPaMZ regimen was included as control in experiments, given its high efficacy in preventing relapse in the RMM and in the Simplici-TB clinical trial (*44*), despite its toxicity, as well as the HRZE standard of care.

The nine experiments were performed across three different laboratories, reflecting the diversity of current experimental models in drug development, including variations in *M. tuberculosis* strains and infection routes. Baseline (pre-treatment) values (median, inter-quartile range) for H37Rv after aerosol infection were 7.78 (7.52-7.78) log10 CFU/lung and 271 (252–271) RS ratio, for H37Rv after intranasal infection 7.43 (7.43-7.56) log10 CFU/lung and 237 (237–272) RS ratio, and for Erdman after aerosol infection 7.33 (7.31-7.33) log10 CFU/lung and 257 (242–257) RS ratio. Incorporating different experimental conditions and their impact on baseline bacterial burden and subsequent relapse prevention is essential for unbiased interpretation of therapeutic efficacy.

### Computational model predicts relapse based on short-term biomarkers, accounting for experimental variability

To predict long-term relapse outcome based on short-term biomarker data, a computational tool was developed and validated using integrated short- and long-term data. An iterative modelling workflow was designed to accommodate the progressively expanding training and validation datasets (**Figure 1**). The initial training dataset 1, consisting of nine unique regimens, contained data on short-term biomarkers, long-term relapse outcome, and relevant baseline and experimental variables. Different modeling techniques were evaluated, and as different machine learning/artificial intelligence (ML/AI) techniques performed similarly to the multivariable logistic regression model, the latter was chosen for continued model development, it being a parsimonious yet powerful modeling technique with easy interpretation and simulation properties. Internal validation was performed using correlation between predicted and observed relapse outcomes, and simulation-based visual predictive checks. Model performance was further assessed using external validation dataset 1, followed by retraining with combined training and validation datasets and subsequent internal and external validation, in a total of three iterations.

After iteration 1, only change of log10 CFU/lung and RS ratio were included as statistically significant predictors, likely due to the smaller dataset size. Between iteration 2 and 3, no difference in the structural model was detected and parameter differences were minor except for the experimental *M. tuberculosis* strain which had a more reasonable estimate for iteration 3 (**Figure 2A, Figure S1**). Final model area under the receiver operator curves (AUROC) values for both the training and validation sets were high for all iterations (0.88-0.93) with an internal AUROC for training and external AUROC for validation datasets in iteration 3 of 0.93 and 0.90, respectively (**Figure 3**). In addition, univariable model development followed by multivariable forward inclusion and backward deletion in iteration 2 and iteration 3 showed similar results (T**ables S2-S5, Figures S2-S3**). Given the minor changes between iterations 2 and 3, and the overlapping estimates for training datasets 2, 3, and validation dataset 3 (**Figure S1**), the confidence in the appropriate model structure and estimates is high.

**Figure 2.**
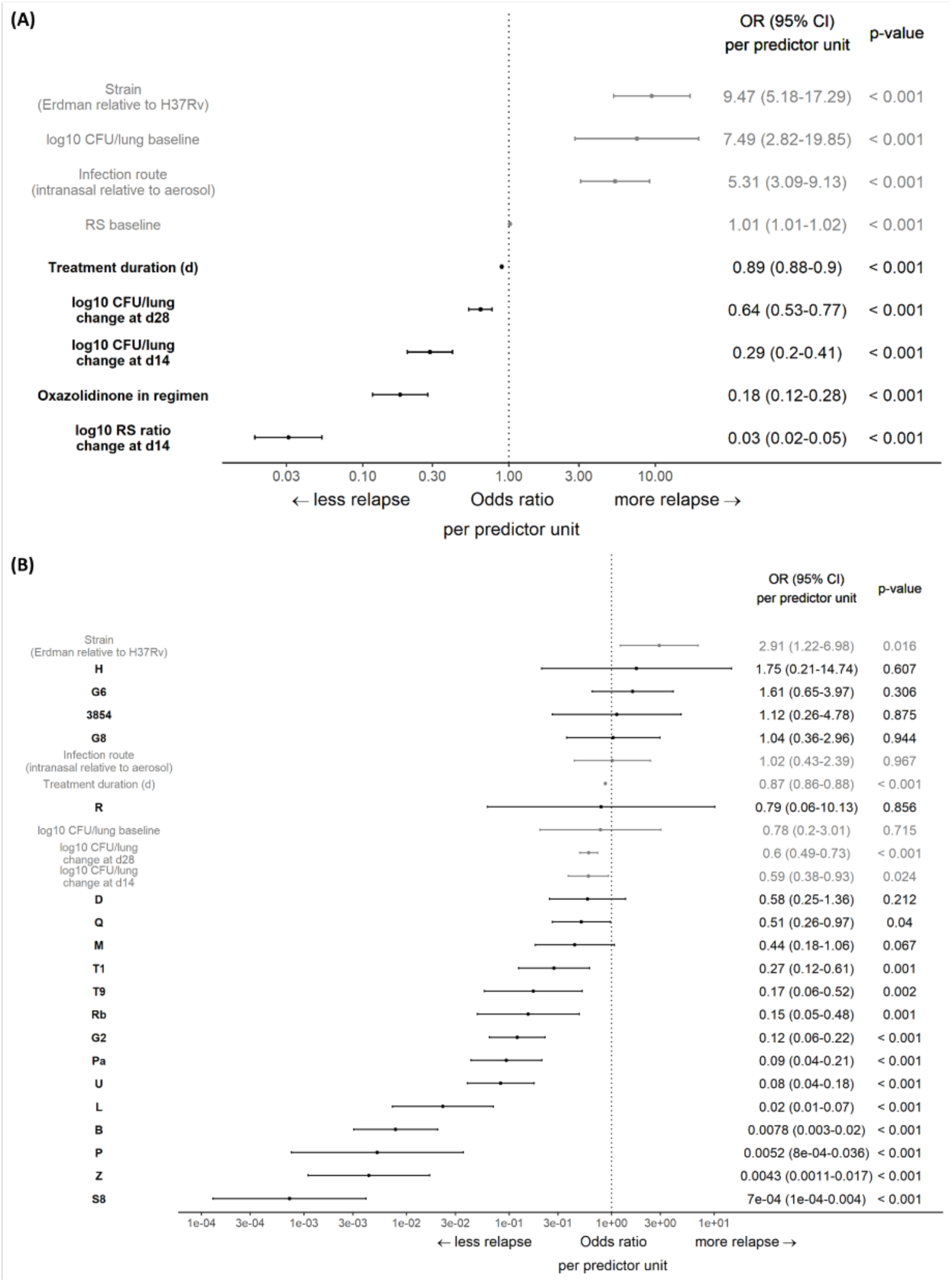
Forest plot of the final model of iteration 3 (**A**) and the model challenged by removing RS ratio biomarker and adding individual sterilizing contribution of drugs to relapse prevention (**B**). Panel shows the final model estimates and their corresponding odds ratio (OR) and 95% confidence interval (CI), for the third and final iteration of the model. Odds ratios represent the association between predictor and outcome, holding other variables constant. Numerical covariates are measured in different scales and comparison is only meaningful when those scales are equivalent. Sterilizing contribution of drugs are at doses used in the experiments modelled here (**Supplementary table 1**). 3854 = MK-3854, B = bedaquiline, CFU = colony forming units, d = days, D = delamanid, G2 = GSK-286, G6 = ganfeborole (GSK-656), G8 = GSK-830, H = isoniazid, L = linezolid, M = moxifloxacin, P = rifapentine, Pa = pretomanid, Q = quabodepistat, R = rifampin, Rb = rifabutin, S8 = sorfequiline (TBAJ-876), T1 = TBD11 (CLB-073), T9 = TBD09 (MK-7762), U = sutezolid, Z = pyrazinamide.

**Figure 3.**
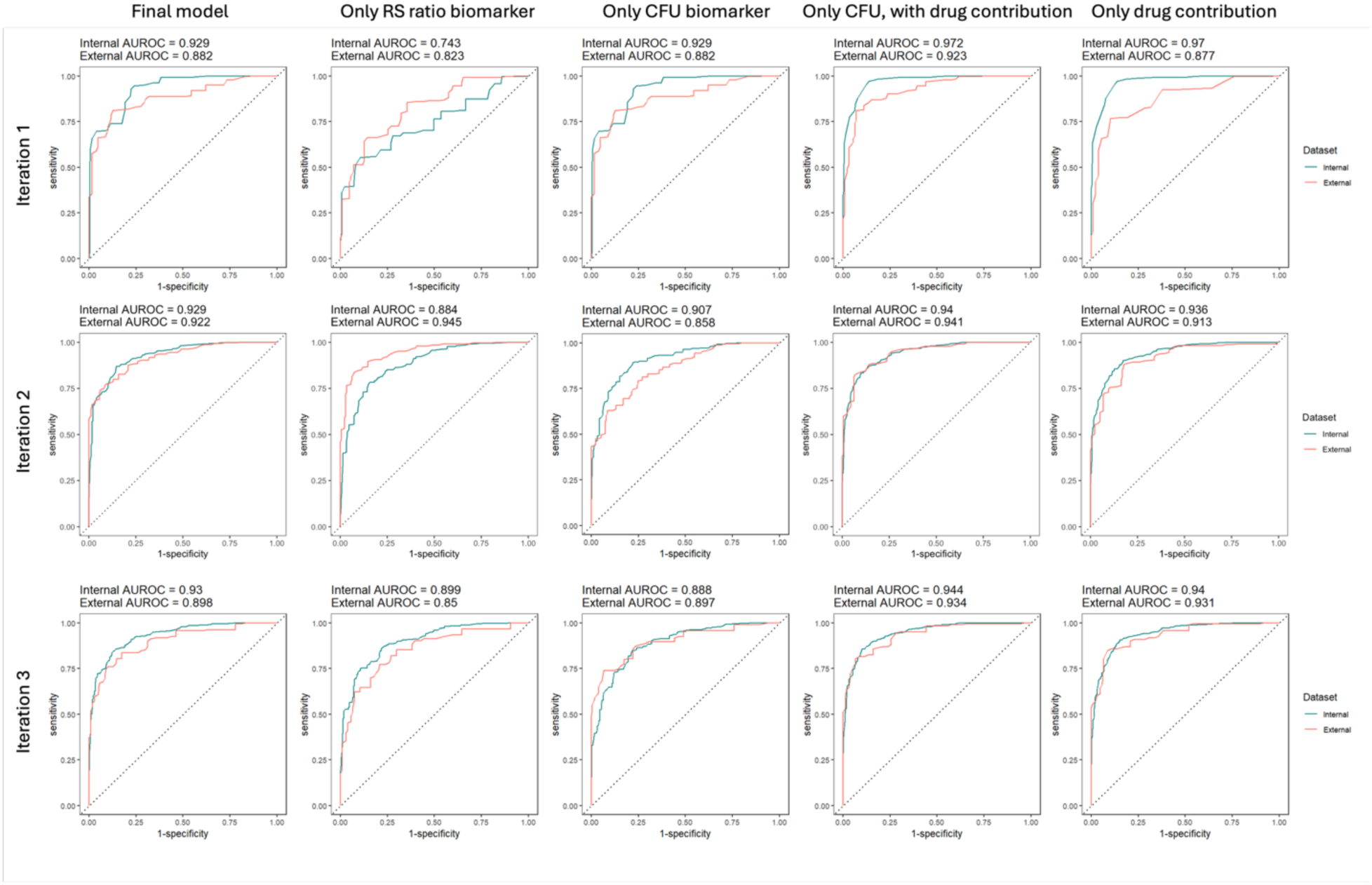
Receiver operator curves (ROC) including their area under the ROC (AUROC) for the iterative model development (**rows**) show no further improvement between iterations 2 and 3 for the final model. Model challenges by removing biomarker predictors (**columns**) show worsened performance, though adding the sterilizing contribution of individual drugs to the relapse prevention restores predictive performance. Training and validation datasets are shown in table 1, for external validation of iteration 2 the contribution of sorfequiline to relapse prevention in the models with drug contribution was imputed by bedaquiline as same drug class, given that no sorfequiline regimens were in training dataset 2.

The final model structure established after iteration 3 included eight predictors, with relapse probability as the dependent variable and treatment duration as the independent variable (**Figure 2**). Four predictors reflected the therapeutic performance on the TB infection and subsequent relapse probability, namely the decrease of log10 CFU/lung from baseline at day 14 and 28, the decrease of RS ratio from baseline at day 14, and the presence of an oxazolidinone in the regimen. Although an interaction term between oxazolidinone presence in the regimen and RS ratio related predictors was tested, based on their mechanism-based interaction, it did not improve model performance. Decrease of RS ratio from baseline at day 28 was statistically significant in the forward inclusion but not maintained in the model after backward deletion. Exploratory data analysis (not shown), suggested that a further decline in log10 CFU/lung between day 14 and 28 could help distinguish regimens, whereas RS ratio changes were less discriminative over that interval. The remaining four predictors reflected experimental conditions, accounting for the *M tuberculosis* strain, route of infection, and baseline values for log10 CFU/lung and RS ratio. Incorporating these variables improved the model’s ability to compare relapse probabilities across experiments and laboratories. Notably, removal of the RS ratio biomarker reduced the model performance significantly in both iterations (**Figure 3**).

The final model was challenged in a sensitivity analysis by assessing the impact of removing either biomarker. Model performance worsened after removing either CFU/lung or RS ratio-related predictors, as expected. The decreased performance after removal of the RS ratio biomarker could be compensated by the inclusion of model-estimated quantitative coefficients for the sterilizing contribution of individual drugs in the regimen (**Figure 2B**). Regimens with drugs known to be sterilizing, such as the diarylquinolines (bedaquiline, sorfequiline), rifamycins (especially rifapentine), and pyrazinamide, showed less relapse for the same change in CFU/lung from baseline than regimens with known non-sterilizing drugs. For drugs in the validation dataset which were not included in the training dataset, the coefficient was assumed to be the same as that of the drug of the same class that was included in the training dataset. A final challenge of the model by only incorporating drug contributions without both CFU/lung and RS ratio biomarkers also performed well (**Figure 3**). These latter two model structures do require prior data from the RMM for all compounds to estimate the quantitative coefficients for their sterilizing contribution.

### Relapse probability over time and time to 95% relapse-free are well predicted for different regimens and experiments

To evaluate the predictive performance of the final model, simulated relapse probability over time accounting for experimental variability was overlaid with the observed proportion of relapsing mice in the experiment (**Figure 4**). For both the 46 regimens in the training dataset and the 14 regimens in the validation dataset, model simulations generally aligned well with the observations. Importantly, the model accurately captured within-regimen differences across experiments (labelled in colors), reflecting the influence of experimental variables and biomarker variability incorporated in the model. For example, BPaQU, BPaL, and BDQU showed slight shifts in observed relapse profiles between experiments, which were also reflected in the model predictions. In the validation set, regimen S8QT9 was not well captured by the predictive model, suggesting a discrepancy between the change in CFU and RS ratio from baseline and the observed relapse for this 3-drug regimen.

**Figure 4.**
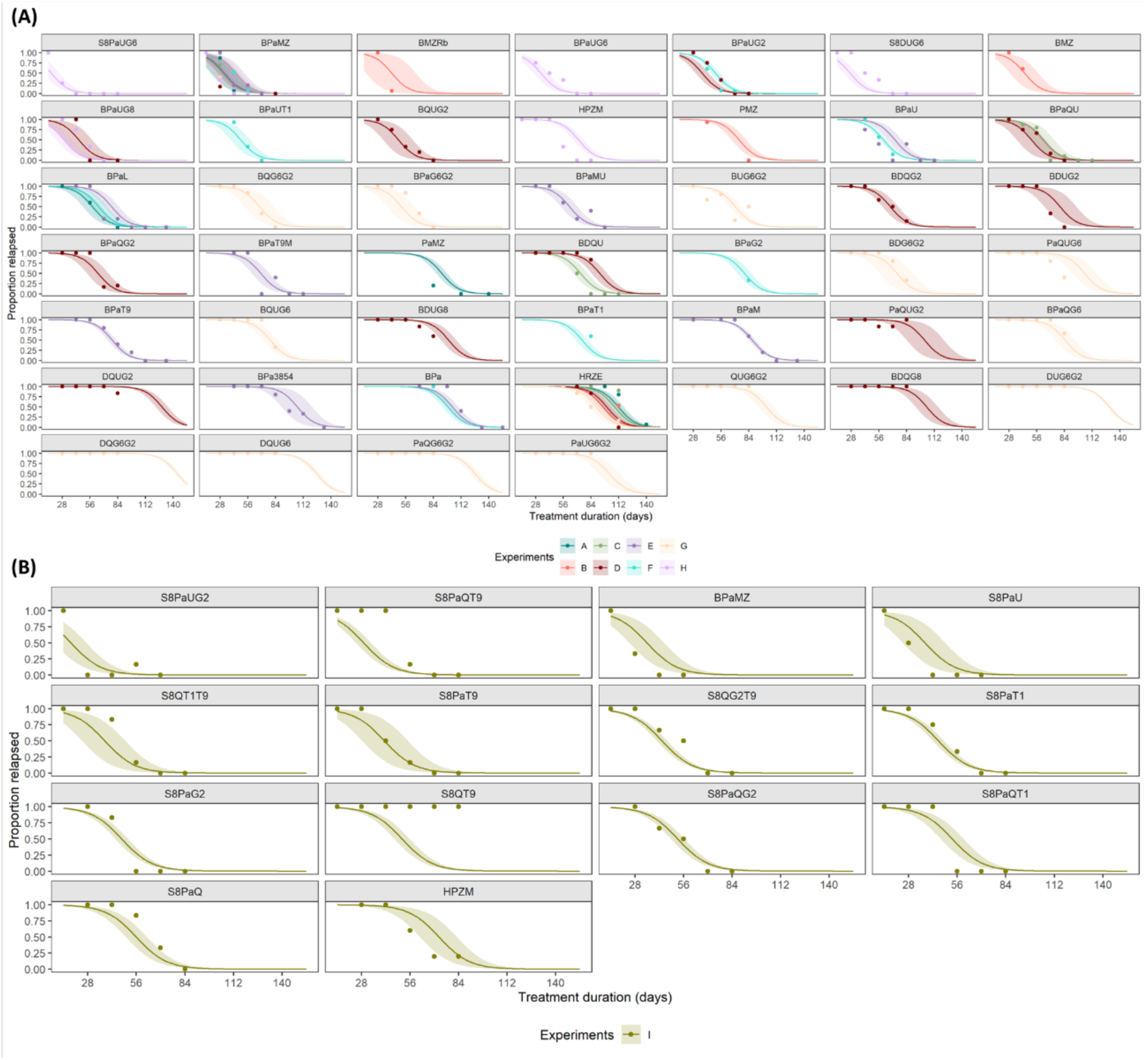
Model-predicted relapse over time of training dataset 3 (**A**) and validation dataset 3 (**B**) by the final model (iteration 3). Observed proportions of relapsing mice are shown in symbols, lines and intervals show median and 95% prediction interval, colors show the experiment. Regimens are defined in Table 1.

Time to 95% relapse-free cure (T95) was determined from the data and the model to further evaluate model performance. Correlation between the observed and predicted T95 was good, with an overall concordance correlation coefficient (CCC, >0.8 considered excellent (*45*)) of 0.73 (95% confidence interval: 0.61-0.81) and a CCC of 0.85 (0.73-0.92) for regimens of interest with T95 values less than or equal to 4 months (**Figure 5**). In the model without RS ratio-related predictors but with the sterilizing contribution of individual drugs to the regimen, the CCC were 0.59 (0.46-0.69) and 0.93 (0.88-0.97), respectively. In the validation dataset, the CCC was 0.67 (0.26-0.88) for the final model, and 0.53 (0.83-0.94) for the model without RS ratio-related predictors but with the sterilizing contribution of individual drugs to the regimen. Based on the predicted T95, regimens could be ranked and showed good agreement with the T95 agnostically determined from the observations (**Figure 6**). For head-to-head comparison even between different experimental settings and baseline values, the model can predict the T95 values while adjusting for experimental variables (**Figure S4-S6**). Consistent with clinical findings (i.e. XBQS trial), regimens combining a diarylquinoline, nitroimidazole, and oxazolidinone with a novel molecule (ganfeborole, GSK-286, GSK-830, or quabodepistat) performed best(*41*). Most of their predicted T95 values were comparable to the BPaMZ control, with S8PaUG6 showing a T95 of 2.5 weeks shorter than BPaMZ. The HPZM regimen, which successfully reduced TB treatment duration from 6 to 4 months in the TBTC S31/A5349 regimen, had a predicted T95 of 86 days, 28 days shorter than the HRZE control. Almost all regimens showing longer T95 values than HRZE lacked a diarylquinoline or included only three drugs. By visualizing the T95 across the short-term biomarker space – defined by log10 CFU/lung at 14 and 28 days, and RS ratio change at 14 days – a visual estimate can be made of the biomarker value which needs to be observed for a target T95 of a novel regimen (**Figure S7**).

**Figure 5.**
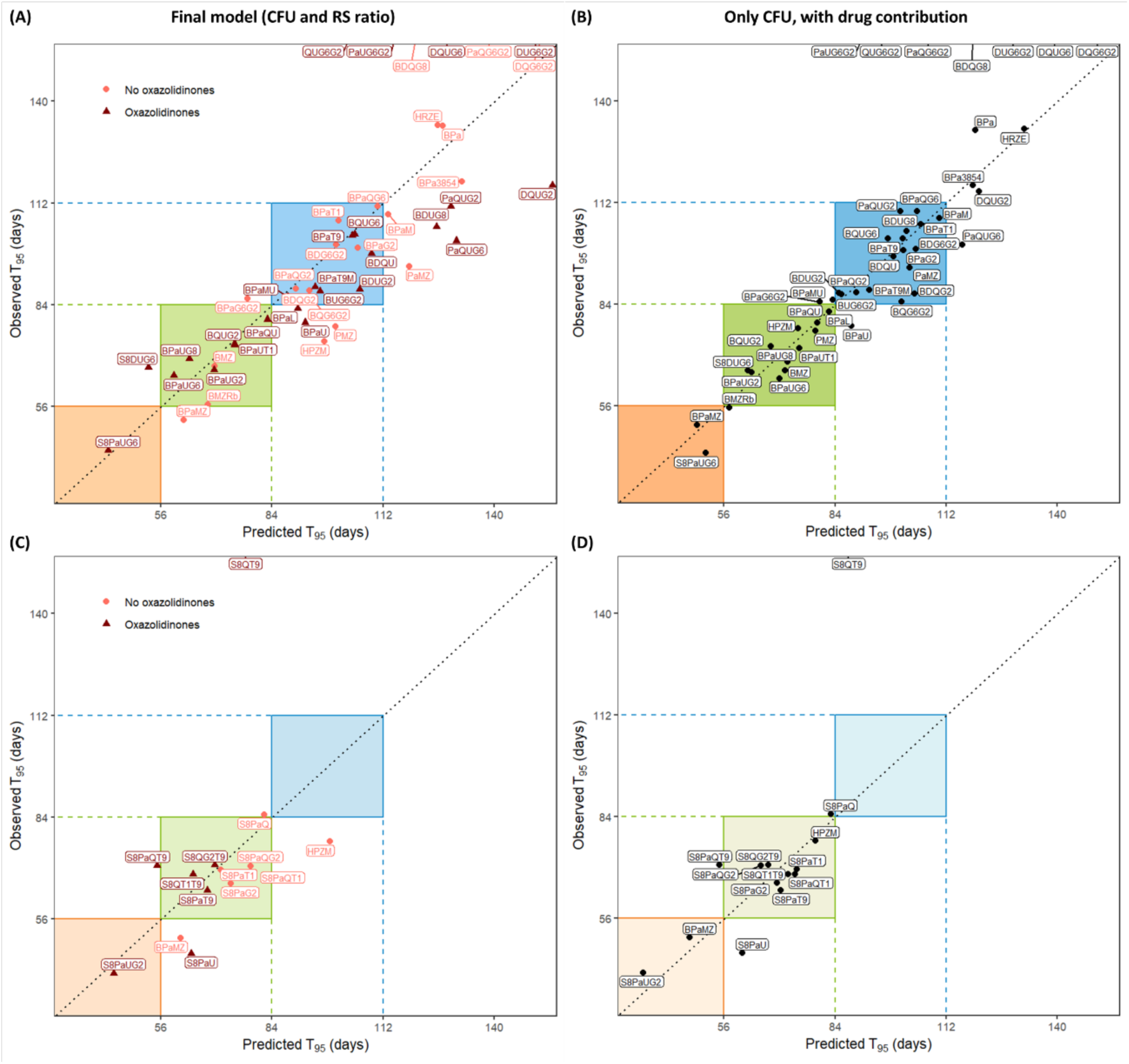
Correlation graph of observed and predicted timepoint at which 95% of mice are relapse-free (T95) for the final model with both biomarkers CFU/lung and RS ratio (**A,C**) and for the model with only CFU/lung biomarker but with the sterilizing drug contribution to relapse prevention (**B,D**) for the iteration 3 training (**top panels A,B**) and validation (**bottom panels C,D**) datasets. Observed T95 was determined by the agnostic fit, predicted T95 was determined by the models (iteration 3). Regimens are defined in Table 1, and predictions are shown here summarized for each regimen across studies, with color and shape depicting if the regimens contain an oxazolidinone (triangle) or not (circle) for panels (**A**) and (**C**).

**Figure 6.**
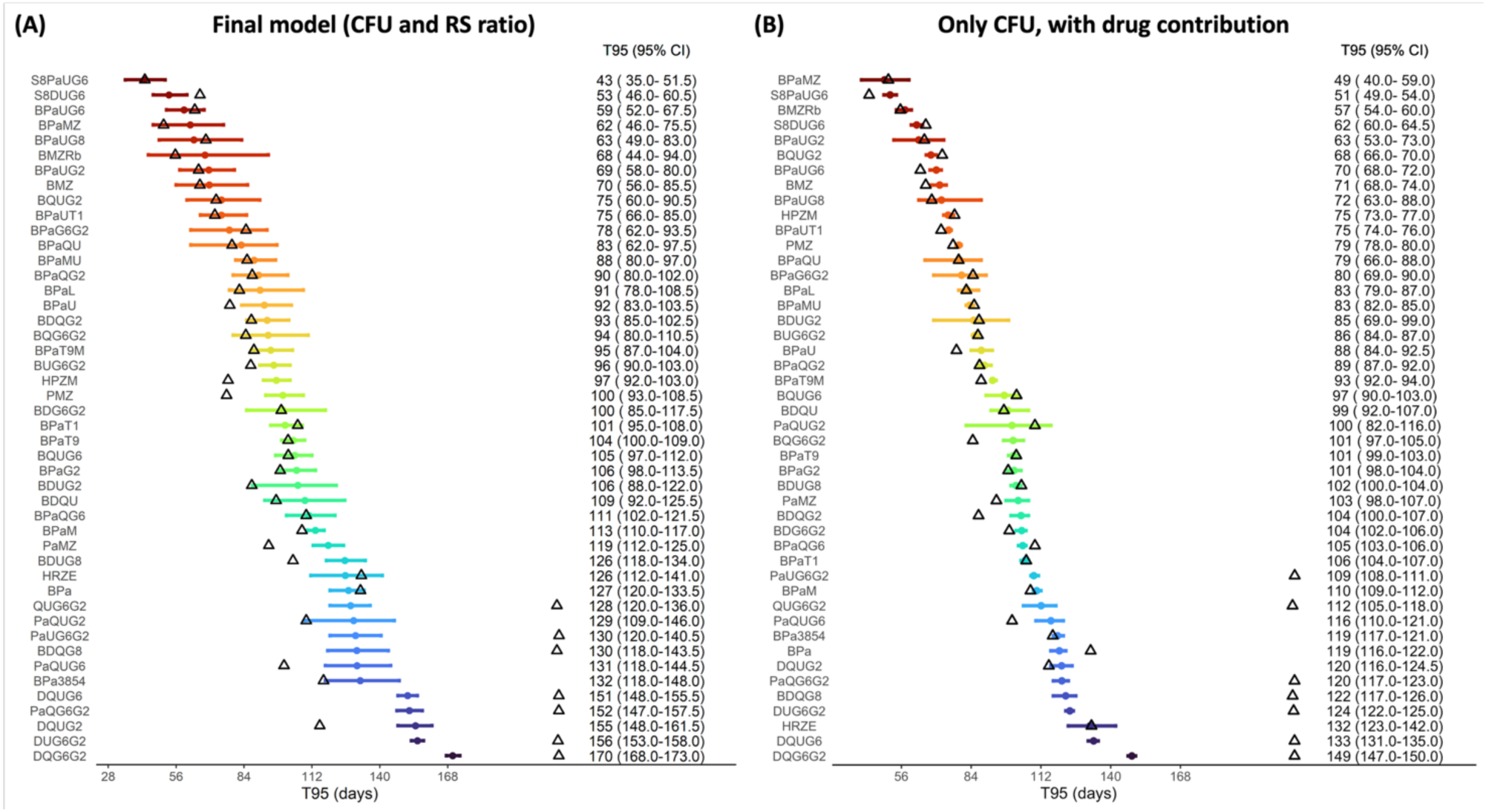
Forest plot of predicted timepoint at which 95% of mice are relapse-free (T95) based on the final model with both biomarkers CFU/lung and RS ratio (**A**) and for the model with only CFU/lung biomarker but with the sterilizing drug contribution to relapse prevention (**B**) for the iteration 3 dataset. Observed T95 was determined by the agnostic fit, predicted T95 was determined by the models (iteration 3). The seven observed T95s at the edge of the graph were from regimens for which no observation was made with less than all mice relapsing. CI = confidence interval, regimens are defined in Table 1.

Regimens were grouped into categories requiring 2, 3, and 4 months of treatment to achieve a 95% relapse-free outcome. This categorical approach was chosen to avoid overinterpreting exact T95 values at this early stage of drug development, where the primary goal remains to shorten treatment to 2-4 months.(*46*) The final model showed high specificity (0.85-0.98) and reasonable sensitivity (0.33-0.70) (**Table 2**), in line with an important purpose of preclinical development: to reliably identify regimens likely to fail in later stages of development. Using the computational model developed here, relapse probability over time, including estimated treatment duration until 95% is relapse-free, can be predicted for novel drug regimens based on innovative early biomarker data, specifically CFU/lung and RS ratio measurements obtained within 4 weeks of treatment, or for regimens of drugs with known sterilizing potential based on CFU/lung and the quantitative sterilizing contribution of the drugs to the regimen.

**Table 2.** Confusion table for final model (iteration 3) performance on the training and validation datasets showing high specificity and reasonable sensitivity by the true positive (TP), false positive (FP), false negative (FN), and true negative (TN) performance as well as the true positive rate (TPR), true negative rate (TNR), positive predictive value (PPV) and negative predictive value (NPV). Care should be taken in interpreting the values for the validation dataset given that only 14 regimens were included, 12 of which were 2- or 3-month regimens.

| *Final model (CFU and RS ratio): training* |  |  |  |  |  |  |  |  |
| --- | --- | --- | --- | --- | --- | --- | --- | --- |
| T95 (mo) | TP | FP | FN | TN | TPR | TNR | PPV | NPV |
| 2 | 1 | 1 | 2 | 42 | 0.33 | 0.98 | 0.50 | 0.95 |
| 3 | 7 | 3 | 5 | 31 | 0.58 | 0.91 | 0.70 | 0.86 |
| 4 | 14 | 4 | 6 | 22 | 0.70 | 0.85 | 0.78 | 0.79 |
| *Only CFU, with drug contribution: training* |  |  |  |  |  |  |  |  |
| T95 (mo) | TP | FP | FN | TN | TPR | TNR | PPV | NPV |
| 2 | 2 | 0 | 1 | 43 | 0.67 | 1.00 | 1.00 | 0.98 |
| 3 | 11 | 3 | 1 | 31 | 0.92 | 0.91 | 0.79 | 0.97 |
| 4 | 17 | 1 | 3 | 25 | 0.85 | 0.96 | 0.94 | 0.89 |
| *Final model (CFU and RS ratio): validation* |  |  |  |  |  |  |  |  |
| T95 (mo) | TP | FP | FN | TN | TPR | TNR | PPV | NPV |
| 2 | 1 | 1 | 2 | 10 | 0.33 | 0.91 | 0.50 | 0.83 |
| 3 | 7 | 4 | 2 | 1 | 0.78 | 0.20 | 0.64 | 0.33 |
| 4 | 0 | 1 | 1 | 12 | 0.00 | 0.92 | 0.00 | 0.92 |
| *Only CFU, with drug contribution: validation* |  |  |  |  |  |  |  |  |
| T95 (mo) | TP | FP | FN | TN | TPR | TNR | PPV | NPV |
| 2 | 2 | 1 | 1 | 10 | 0.67 | 0.91 | 0.67 | 0.91 |
| 3 | 8 | 2 | 1 | 3 | 0.89 | 0.60 | 0.80 | 0.75 |
| 4 | 0 | 1 | 1 | 12 | 0.00 | 0.92 | 0.00 | 0.92 |

## DISCUSSION

Based on an integrated dataset of nine RMM experiments and three iterations of model training and external validation, a computational model was developed to successfully predict long-term relapse probability in mice over time using short-term biomarkers, specifically CFU and the innovative RS ratio biomarker. Across iterations, the structural model remained consistent, with only minor changes in predictor coefficient estimates between the second and third iterations.

External validation confirmed the model’s robustness. Challenging the model to assess removal of either biomarker, showed that the predictive model with CFU/lung biomarker but without RS ratio performed similarly to the final model when adjusting for the sterilizing contribution of individual drugs in the regimen. The latter, however, requires long-term outcome data, instead of the short-term biomarker data, and would be less suitable for novel compounds. By quantifying both therapeutic and experimental variables, the model predictions account for experimental variability, enabling unbiased comparison of regimen performance. This approach allows for early ranking of novel regimens based on data from just 4 weeks of treatment in mice. As a result, follow-up experiments can be strategically prioritized, preventing unnecessary lengthy, resource-demanding experiments requiring larger numbers of mice to evaluate relapse prevention and focusing efforts only on the regimens with most potential.

Results of the BALB/c RMM are informative of clinical efficacy. The standard of care HRZE, developed more than 40 years ago, exhibited a long predicted T95 exceeding 4 months. In contrast, HPZM, the first regimen to successfully shorten treatment from 6 to 4 months in clinical trials (*3, 47*), had an intermediate T95 of approximately 3 months. BPaMZ, the most clinically efficacious regimen tested to date (though discontinued due to safety concerns (*44*)) showed a T95 of less than 2 months. Based on the integrated data and the iteratively developed computational model, the regimens BDQU and BPaQU, currently under evaluation in the Gates MRI-TBD06-201 PAN-TB trial (i.e. XBQS), are predicted to perform similar to, and approximately 3 weeks better than, HPZM, respectively, in terms of relapse prevention in mice. This aligns with BPaQU but not BDQU showing similar performance after 4 months of treatment compared to 6 months of standard of care at the 12-month trial observation(*41–43*).

Additionally, the S8PaL regimen, under evaluation in the TB Alliance NC-009 trial, is expected to perform comparably to BPaMZ, assuming similar efficacy for the oxazolidinones sutezolid and linezolid. The complexity of translating BALB/c RMM results into clinical success is exemplified by clofazimine-containing regimens (*48*). Despite promising outcomes in preclinical (BALB/c) studies, these regimens failed to demonstrate efficacy in the CLO-FAST clinical trial (*49*). The discrepancy underscores the importance of incorporating additional pharmacological dimensions into regimen evaluation. One important consideration if systemic pharmacokinetics for regimens, possible drug-drug interactions, and predicting those from preclinical species to patients. Another critical factor is target site exposure, which has emerged as a hallmark of pharmacological success (*50, 51*). Integrating this parameter into computational modelling frameworks offers a more holistic approach to regimen selection, enhancing predictive accuracy and translational relevance (*19, 38*).

It is known that changes in CFU/lung from baseline cannot always distinguish sterilizing from non-sterilizing regimens (*24–26*). For regimens consisting of drugs with known sterilizing potential, such as diarylquinolines, rifamycins, and pyrazinamide, the addition of a quantitative coefficient for known drugs to the model sufficed for good predictive performance of relapse prevention. However, such a model needs relapsing mouse data to estimate the coefficients for the sterilizing contribution of the individual drugs (*25, 26*). As such, for novel drugs currently in the pipeline, the short-term RS ratio biomarker can inform on the sterilizing potential as reflected by the drug’s impact on the physiological state of the mycobacteria. For those regimens with novel compounds that are prioritized based on the short-term RS ratio and CFU/lung biomarker data, confirmatory experiments in the relapsing mouse model can be performed. These data can feed back into the model to quantify the sterilizing contribution of those novel drugs and facilitate additional regimen predictions.

The developed model accounts for experimental variability both within and between experiments and laboratories, enabling unbiased comparisons of regimen performance. Importantly, the integrated dataset includes two of the most frequently used strains of *M. tuberculosis* and two of the most frequently applied routes of administration. As a result, the model is broadly applicable and likely to generalize well across a wide range of global preclinical studies. Nevertheless, further evaluation and validation of other experimental conditions or more diverse *M. tuberculosis* strains, including other phylogenetic lineages, is warranted to expand its applicability (*18*). At this stage, the dose was not explicitly incorporated into the model as most drugs were tested at a single standardized dose (**Table S1**). However, as the short-term biomarkers CFU/lung and RS ratio are predictors rather than dose, it is expected that dose variations may inherently be captured through their response of these biomarkers. That is, a higher dose resulting in a higher decrease in CFU/lung would impact the relapse prediction through the change in CFU/lung, as for example visible for BPaQU (**Figure 3**). Another advantage is that, when informing the predictive model, the ultra short-course mouse model of 4 weeks of treatment can be performed without full understanding of the drug’s sterilizing potential or the dose-exposure-response relationship *in vivo*, given that the model predicts a T95 based on the biomarker response to the given dose(s). This is particularly advantageous during early drug development when dose-exposure-response relationships may still be under investigation. The outcome of this 4-week treatment experiment, informed by the computational model developed here (**Figure S7**), can guide prioritization of regimens and inform the design of subsequent studies to characterize the PKPD relationships. This biomarker-driven approach contrasts with other relapse prediction models that rely on regimen composition to estimate relapse probability (*20, 25, 26, 42, 43*). A key advantage of the short-term biomarker driven approach is the speed at which predictions can be generated, requiring only 4 weeks of experimental treatment, and substantially less mice compared to the full relapsing mouse model. Concretely, for the short-term biomarkers necessary to develop our predictive model, only approximately one-third of mice were necessary, showing two-third reduction in experimental animal use. However, compound and regimen-informed models remain valuable for *de novo* regimen performance predictions, as long as relapse data is available on regimens containing the drugs of interest. This approach can also characterize the dose-dependent sterilizing potential of compounds based on dose-ranging RMM data, complementary to the biomarker approach with dose implicitly captured through dose-dependent biomarker response (*25, 26*).

The work presented here has several limitations. The predictive model assumes a common slope across all regimens, implying that the rate at which relapse probability declines with increasing treatment duration is the same for all regimens, and this relationship can only shift left or right. A sensitivity analysis allowing for regimen-specific slopes yielded very similar results for almost all regimens, consistent with previous findings comparing the slopes of HRZE, HPZM, HRZM, BPaMZ, and BPaL (*18, 52*). Moreover, inclusion of different slopes per regimen will diminish the model’s forecasting performance, as it would need relapse data for each regimen to estimate the slope parameter (*18, 52*).

Another limitation is the model’s reliance on RS ratio as a key predictor. While this biomarker significantly improves model performance (**Figure 3**) and responds more rapidly than other mycobacterial biomarkers, its interpretation becomes more complex when regimens include oxazolidinones (*32*). To address this, a binary predictor for the presence of oxazolidinones was included *a priori*. However, further refinement by estimating a statistical interaction term between oxazolidinones in the regimen and RS ratio related predictors did not improve model performance. Thus, while the RS ratio remains a valuable and statistically significant predictor, results involving oxazolidinones or protein synthesis inhibitors should be interpreted with caution. The three-drug regimen S8QT9 in the validation dataset was not captured well by the predictive model which was mainly developed based on four-drug regimen data, suggesting careful interpretation for regimens with less than four drugs. Finally, logistic regression was selected as the most appropriate modelling approach given the current dataset size. However, as the dataset size continues to grow, the potential application of ML/AI techniques should be re-evaluated (*53*).

In conclusion, the computational model developed in this study enables the prediction of relapse probability based on data from a 4-week treatment experiment. This approach allows for early prioritization of regimens with the highest potential for sterilizing activity, guiding the selection of candidates for confirmatory relapsing mouse model studies and subsequent clinical evaluation. By streamlining preclinical decision-making, the model supports more efficient and targeted TB drug development.

## MATERIALS AND METHODS

### Study design

The objective of this study was to develop a computational model to rank regimens by their relapse prevention potential on the long-term, informed by innovative biomarker RS ratio and CFU/lung on the short-term. To this objective, the predictors of relapse were quantified in an iterative model development and validation approach, the impact of experimental variables related to the experimental design and inter-experimental variability were distinguished from therapeutic variables of interest, and as a result regimens were ranked objectively accounting for these experimental sources of variability for an unbiased comparison.

Data from 9 published and unpublished experimental studies were integrated for model development and validation (*42, 43*). Predictors included short-term biomarker observations after 14 and 28 days of treatment for CFU/lung and RS ratio, while the primary outcome was relapse probability and the derived time to 95% relapse-free cure (T95), with relapse-free cure defined as 0 CFU/lung 12 weeks after termination of treatment. In addition, experimental variables including baseline values, and variables on the composition of the regimen were included. Different mathematical modelling techniques were employed to characterize the probability of relapse over time based on these predictors. Model evaluation was performed based on simulation-based visual predictive checks, concordance correlation coefficient, specificity and sensitivity, and the area under the receiver operator curve (AUROC). The model was iteratively trained and validated on the data from the 9 experimental studies becoming available over time.

### Mouse experiments for short-term biomarkers and relapse outcome

All animal experiments were approved by the relevant institutional ethical committee. At Johns Hopkins University, all housing and procedures involving mice were approved by the Animal Care and Use Committee at Johns Hopkins University School of Medicine (JHU). JHU’s animal welfare assurance number is A3272.01. JHU is registered with the USDA to conduct animal research and has maintained active AAALAC accreditation since 10/4/1974. At Colorado State University, all procedures and protocols for infecting mice with M. tuberculosis (Mtb) and subsequent drug treatments in the described mouse infection studies were approved by the Colorado State University Institutional Animal Care and Use Committee (IACUC) (Reference numbers: 17-7701A and 18-7912A). CSU IACUC accreditation: Association for Assessment and Accreditation of Laboratory Animal Care International (CSU Unit 000834). The Evotec France SAS animal facility is accredited by the French Ministry of Agriculture and by the Association for Assessment and Accreditation of Laboratory Animal Care International (AAALAC). All studies were performed under the European Communities Council Directive (2010/063/EU) for the care and use of laboratory animals and approved by local Ethical Committee CEPAL: CE 029 and authorized by the French Ministry of Education, Advanced Studies, and Research.

In short, female BALB/c mice were infected with *M. tuberculosis*, either H37Rv (American Type Culture Collection strain ATCC 27294) or Erdman (ATCC 35801) strain, through high dose aerosol or intranasal routes of infection, depending on which laboratory performed the experiment (**Table 1**). Mice were randomly allocated to treatment, which started at 14 days post-infection and was comprised of daily dosing by gavage administration, 5 out of 7 days per week. Details on drug dose and manufacturer per regimen are shown in **Table S1**. Mice were sacrificed after infection, at start of treatment (defined as baseline), and at 2, 4 and 8 weeks after start of treatment to quantify bacterial burden (colony forming units [CFU] per lung normalized to the entire lung mass) by quantitative cultures on complete Middlebrook 7H11 medium (Becton Dickinson) supplemented with glycerol (Fisher), 10% (v/v) acid-albumin-dextrose-catalase (OADC, GIBCO), 0.4% (w/v) activated charcoal (JT Baker), and cycloheximide [10 mg/mL], carbenicillin [50 mg/mL], polymixin B [25 mg/mL] and trimethoprim [20 mg/mL] (all 4 antibiotics from Sigma). RS ratio (*28, 29, 32*) was quantified by a QX100 Droplet Digital PCR system (Bio-Rad) from lung homogenates. After completion of treatment, a 3-month wash-out period was observed after which mice were sacrificed to determine relapse or no relapse as defined by the presence or absence, respectively, of observable CFU/lung by quantitative culture. For relapse timepoints measured at CSU or JHU, 7H11-OADC lacking activated charcoal was used, while Evotec utilized 7H11-OADC + 0.4% activated charcoal.

### Data consolidation

Iterative dataset compilation into training and validation sets was performed, as available datasets were used as training datasets and new datasets once available served as external validation sets, after which the combined datasets served as training datasets for iterative model development and validation (**Table 1**). Training dataset 1 contained datasets A, B, C, with 167 short-term biomarker observations of CFU/lung and RS ratio, and 497 long-term relapse observations, while validation dataset 1 contained datasets D with 144 short-term biomarker observations of CFU/lung and RS ratio, and 349 long-term relapse observations. In the next iteration, training dataset 2 contained datasets A-F with an additional 332 short-term biomarker observations of CFU/lung and RS ratio, and 815 long-term relapse observations, compared to training dataset 1. Validation dataset 2 contained datasets G and H with 199 short-term biomarker observations of CFU/lung and RS ratio, and 532 long-term relapse observations. In the last iteration, training dataset 3 contained datasets A-H with an additional 199 short-term biomarker observations of CFU/lung and RS ratio, and 532 long-term relapse observations compared to training dataset 2. Validation dataset 3 contained dataset I with 145 short-term biomarker observations of CFU/lung and RS ratio, and 395 long-term relapse observations.

Primary binary outcome of relapse at the designated treatment duration was linked to the corresponding continuous experimental (baseline log10 CFU/lung and RS ratio, *M. tuberculosis* strain, administration route, exact washout duration) and therapeutic efficacy (CFU/lung and RS ratio at 2 and 4 weeks, change in CFU/lung and RS ratio at 2 and 4 weeks from baseline, change in CFU/lung and RS ratio from 2 to 4 weeks, as well as presence or absence of oxazolidinones in the regimen) data, and standard deviations for all continuous variables were determined (**Table S2-3**).

### Model development

To quantify the relationships between short-term predictors and long-term relapse outcome in the mouse data, different mathematical modelling techniques were employed. These techniques included statistical regression methods such as logistic regression, as well as machine learning/artificial intelligence (ML/AI) techniques such as decision trees, random forests, and neural networks. Testing ML/AI techniques was performed to assess improved accuracy over more standard logistic regression modelling, which has the advantage of interpretability and simulations of different relapse scenarios based on the quantified predictors and variability. Logistic regression model development followed the standard approach of univariable analysis followed by stepwise multivariable analysis with forward inclusion until no further predictor improved the model statistically significantly (p<0.05) after which predictors were backward deleted until no further predictor could be removed without statistical significance (p>0.01). The presence of an oxazolidinone in the regimen was included as a predictor *a priori*, because this drug class stabilizes short-lived rRNA resulting in the RS ratio decreasing less, as the rate of decline of the numerator of the ratio is delayed by this stabilization. As the purpose of the model was to predict the relapse probability of novel regimens with or without an oxazolidinone or other translational inhibitor, it was important to account for this interaction between the biomarker and drug class. In addition, an interaction term between the presence of oxazolidinones and the RS ratio based predictors was tested. Model selection was based on specificity and sensitivity, and AUROC. The final model for each iteration was challenged in a sensitivity analysis to assess the impact of either CFU/lung or RS ratio biomarker by removal of those biomarker-related predictors from the final model followed by re-estimating the coefficients and running diagnostics. In addition, the contribution of individual drugs to the regimen’s relapse prevention potential was tested by estimating coefficients for each drug in the training dataset that could be distinguished from the remainder of the regimen. In case of regimens in the external validation containing drugs that were not part of the training dataset, the coefficient was imputed from the same drug class.

To determine the specificity and sensitivity, the observed T95 needed to be determined, which was done through agnostic logistic regression modelling specific to that regimen in that experiment, and informed by the relapse data, with the purpose of deriving an agnostic T95 value to serve as a ground truth, given that an observation of that exact nature is not common within the experimental constraints.

Both agnostic and model-predicted T95 values were calculated based on the probability of relapse:

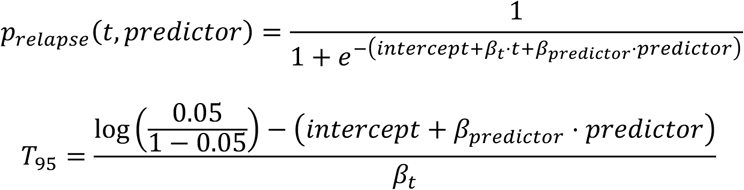

where the probability of relapsing, *p_relapse_* is a function of predictors and their coefficient (*β_predictor_*), as well as treatment duration (*t*) as independent variable, as longer treatment duration will lower relapse probability irrespective of the regimen. For the agnostic T95, the coefficient was estimated separately for each regimen in each experiment, while for the predictive model, the coefficients were estimated for all predictors that were found statistically significant to determine the magnitude of the relapse probability being impacted by the predictor. Estimates of the coefficient can be compared to each other when the predictor unit are on the same scale (e.g. log10 CFU/lung). This modeling approach is based on the assumption that a single slope (*β_t_*) shifts with different predictor values, an assumption which was tested in a sensitivity analysis and has been shown to be valid for datasets of the sizes relevant here.

### Model diagnostic and validation

The iterative workflow included internal and external validation in 3 iterations (**Table 1**). Internal validation to diagnose model performance was performed on the corresponding training dataset, but given the limited number of regimens especially in the first training datasets (9, 29, or 46) no training:testing cut could be performed. More importantly, external validation was performed for each of the 3 iterations (**Table 1**), which was performed after completion of each model development iteration, and blinded to the model building. Model diagnosis and internal and external validation were performed based on numerical, graphical, and pharmacological criteria. Numerically, assessment was based on the AUROC as well as on the specificity (true negative rate) and sensitivity (true positive rate) for categorizing regimens as 2, 3, or 4 month regimens, and the concordance correlation coefficient to determine agreement between observed and predicted T95 values along the line of identity (*y* = *x*). Graphically, simulation-based visual predictive checks were performed in which for each regimen within each experiment, 500 simulations were performed including stochastic variability in CFU/lung and RS ratio biomarkers through a normal distribution with mean and standard deviation as determined in the experimental dataset corresponding to that regimen. The model predictions for these 500 simulations were summarized as median (line) and prediction intervals of 2.5th and 97.5th quantiles (area) over time, and overlaid with the observed proportion of relapsing mice. In addition, receiver-operator curves (ROCs) as well as correlation graphs of observed and predicted T95 values were assessed. Pharmacological knowledge and understanding about the relationship between microbiological, therapeutic, and experimental predictors were employed for model outcome assessment.

### Regimen ranking based on experimental variables

Ranking regimens within an experiment can be performed without correcting for experimental variables such as baseline CFU/lung and RS ratio, *M. tuberculosis* strain, and route of administration. However, to compare regimen performance between experiments, both within and between laboratories, regimen ranking needs to account for these experimental variables. To that aim, for each combination of *M. tuberculosis* strain and route of administration, n=500 datasets were simulated with median CFU/lung and RS ratio baseline values and CFU/lung and RS ratio values sampled from a normal distribution of their mean and standard deviation determined in the experimental dataset corresponding to that regimen.

Ranking was performed based on calculated T95 and visualized with the 2.5th and 97.5th quantile of the T95 as determined from the simulation.

### Software and statistical analysis

This meta-analysis of 9 datasets was performed, where model development was performed in a stepwise selection approach with forward inclusion (p<0.05) and backward elimination (p>0.10) to identify statistically significant predictors. Statistical testing was performed through the likelihood ratio test, assuming Χ^2^-distribution and a decrease of 3.84 points equaling p=0.05 for 1 degree of freedom. No random split in training and testing datasets was performed given the limited number of regimens, but external validation was performed for each of the three iterations with data of 14, 20, and 14 different regimens, respectively, blinded to the model development process of the model being validated. Model-based simulations were performed n=500 times based on sampling from a normal distribution reflecting variability in the original data. All data handling, analysis, and visualizations were performed in R (version 4.4.1) through the RStudio interface (version 2023.09.0), with packages tidyverse (version 2.0.0) including dplyr (version 1.1.4), tidyr (version 1.3.1), and ggplot2 (version 3.5.1), and pROC (version 1.18.5) (*54*).

## List of Supplementary Materials

Figures S1 to S7

Tables S1 to S5

## Acknowledgments

The authors gratefully acknowledge the contributions of the Project to Accelerate New Treatments for Tuberculosis (PAN-TB) Consortium funded by the Gates Foundation and its Consortium members, specifically Evotec as data generator, and all asset owners including Gates Medical Research Institute, GSK, Johnson & Johnson, Otsuka Pharmaceutical, and TB Alliance. The authors gratefully acknowledge the support from the National Institutes of Health (NIH) of the Preclinical Design and Clinical Translation of TB Regimens (PReDicTR) Consortium under award number UM1AI179699, funding from the Gates Foundation under grant ID numbers INV-009105, ‘TB Drug Accelerator: TB mouse in vivo models’ [GTR, NDW], INV-009549, ‘Validation of Mtb rRNA as a PD Marker of Treatment Response’ [NDW, GTR, RMS], INV-008993 ‘Identify the best TB drug regimen candidates through non-clinical evaluation’ [SS, AMU], and funding from the Gates Medical Research Institute under investment ID numbers 56315 and 57159 [ELN]. The content is solely the responsibility of the authors and does not necessarily represent the official views of the NIH, and the conclusions and opinions expressed in this work are those of the author(s) alone and shall not be attributed to the Gates Foundation. Merck & Co., Inc., USA (known as MSD outside the United States and Canada) is acknowledged as asset owner for TBD09. Hoang-Anh Vu is acknowledged for his valuable assistance. Figure 1 was created in BioRender. Savic, R. (2025) https://BioRender.com/a3epvem.

## Funding

National Institutes of Health grant UM1AI179699 (RCvW, BPS, LC, ELN, GTR, NDW, RMS)

Gates Foundation grant INV-009105 (GTR, NDW)

Gates Foundation grant INV-009549 (GTR, NDW, RMS)

Gates Foundation grant INV-009105 (GTR)

Gates Foundation grant INV-008993 (SS, AMU)

Gates Medical Research Institute grant ID-56315 (ELN)

Gates Medical Research Institute grant ID-57159 (ELN)

U.S. Centers for Disease Control and Prevention (Westat subcontract 8758-S01) (ELN)

Veterans Affairs 1I01BX004527-01A1 (NDW)

## Author contributions

Conceptualization: ELN, GTR, NDW, RMS

Methodology: RCvW, BPS, LC, SS, AMU, ELN, GTR, NDW, RMS

Investigation: RCvW, BPS, LC, SS, ELN, GTR, NDW Visualization: RCvW, BPS, LC

Funding acquisition: AMU, ELN, GTR, NDW, RMS Project administration: RCvW, RMS

Supervision: AMU, ELN, GTR, NDW, RMS Writing – original draft: RCvW, BPS, LC

Writing – review & editing: RCvW, BPS, LC, SS, AMU, ELN, GTR, NDW, RMS

## Data and materials availability

The study data is publicly available through the Critical Path Institute; TB-Platform for the Aggregation of Preclinical Experiments Data (TB-APEX) database and in original publications (details in Table S1), as well as with the final model code at Zenodo (10.5281/zenodo.21397999).

