## Supplementary Materials for "Predicting tuberculosis relapse based on 28-day CFU, RS ratio, and/or drug contribution for novel regimens in the relapsing mouse model"

**Table S1.** Regimens and doses tested in the BALB/c relapsing mouse model in the included datasets. [i] training set 1; [ii] validation set 1; [iii] training set 2; [iv] validation set 2; [v] training set 3; [vi] validation set 3

| Dataset | Data source | Regimen | Abbr. |
|---|---|---|---|
| A [i,iii,v] | Walter ND et al, Nat Commun. 2021 12(1):2899. doi: 10.1038/s41467-021-22833-6 | Isoniazid (10 mg/kg)<br>Rifampin (10 mg/kg)<br>Pyrazinamide (150 mg/kg)<br>Ethambutol (100 mg/kg) | HRZE |
| | | Pretomanid (100 mg/kg)<br>Moxifloxacin (100 mg/kg)<br>Pyrazinamide (150 mg/kg) | PaMZ |
| | | Bedaquiline (25 mg/kg)<br>Pretomanid (100 mg/kg)<br>Linezolid (100 mg/kg) | BPaL |
| | | Bedaquiline (25 mg/kg)<br>Pretomanid (100 mg/kg)<br>Moxifloxacin (100 mg/kg)<br>Pyrazinamide (150 mg/kg) | BPaMZ |
| B [i,iii,v] | Tasneen R et al,<br>Antimicrob Agents Chemother. 2022 66(4):e0239821.<br>doi: 10.1128/aac.02398-21<br><br>Dide-Agossou C et al,<br>Antimicrob Agents Chemother. 2022 66(4):e0231021.<br>doi: 10.1128/aac.02310-21 | Isoniazid (10 mg/kg)<br>Rifampin (10 mg/kg)<br>Pyrazinamide (150 mg/kg)<br>Ethambutol (100 mg/kg) | HRZE |
| | | Rifapentine (10 mg/kg)<br>Moxifloxacin (100 mg/kg)<br>Pyrazinamide (150 mg/kg) | PMZ |
| | | Bedaquiline (25 mg/kg)<br>Moxifloxacin (100 mg/kg)<br>Pyrazinamide (150 mg/kg) | BMZ |
| | | Bedaquiline (25 mg/kg)<br>Moxifloxacin (100 mg/kg)<br>Pyrazinamide (150 mg/kg)<br>Rifabutin (5 mg/kg) | BMZRb |
| C [i,iii,v] | Primary manuscript in submission<br><br>Data available at 10.5281/zenodo.21397999 | Isoniazid (10 mg/kg)<br>Rifampin (10 mg/kg)<br>Pyrazinamide (150 mg/kg)<br>Ethambutol (100 mg/kg) | HRZE |
| | | Bedaquiline (25 mg/kg)<br>Delamanid (6 mg/kg)<br>Quabodepistat (9 mg/kg)<br>Sutezolid (50 mg/kg) | BDQU |
| | | Bedaquiline (25 mg/kg)<br>Pretomanid (50 mg/kg)<br>Quabodepistat (9 mg/kg)<br>Sutezolid (50 mg/kg) | BPaQU |
| | | Bedaquiline (25 mg/kg)<br>Pretomanid (100 mg/kg)<br>Moxifloxacin (100 mg/kg)<br>Pyrazinamide (150 mg/kg) | BPaMZ |
| D [ii,iii,v] | Critical Path Institute;<br>TB-Platform for the Aggregation of Preclinical Experiments Data (TB-APEX) Wave 2022, EV-TL-TBa22001 | Bedaquiline (25 mg/kg)<br>Delamanid (6 mg/kg)<br>Quabodepistat (9 mg/kg)<br>Sutezolid (50 mg/kg) | BDQU |
| | | Bedaquiline (25 mg/kg)<br>Pretomanid (50 mg/kg)<br>Quabodepistat (9 mg/kg)<br>Sutezolid (50 mg/kg) | BPaQU |
| | | Bedaquiline (25 mg/kg)<br>Pretomanid (100 mg/kg)<br>Moxifloxacin (100 mg/kg)<br>Pyrazinamide (150 mg/kg) | BPaMZ |
| | | Bedaquiline (25 mg/kg)<br>Delamanid (2 mg/kg)<br>Quabodepistat (14 mg/kg)<br>GSK-830 (5 mg/kg) | BDQG8 |
| | | Bedaquiline (25 mg/kg)<br>Delamanid (2 mg/kg) | BDQU |

| Dataset | Data source | Regimen | Abbr. |
|---|---|---|---|
| | | Quabodepistat (14 mg/kg)<br>Sutezolid (100 mg/kg) | |
| | | Bedaquiline (25 mg/kg)<br>Delamanid (2 mg/kg)<br>Sutezolid (100 mg/kg)<br>GSK-286 (35 mg/kg) | BDUG2 |
| | | Bedaquiline (25 mg/kg)<br>Pretomanid (40 mg/kg)<br>Sutezolid (100 mg/kg)<br>GSK-286 (35 mg/kg) | BPaUG2 |
| | | Bedaquiline (25 mg/kg)<br>Delamanid (2 mg/kg)<br>Sutezolid (100 mg/kg)<br>GSK-830 (5 mg/kg) | BDUG8 |
| | | Bedaquiline (25 mg/kg)<br>Pretomanid (40 mg/kg)<br>Sutezolid (100 mg/kg)<br>GSK-830 (5 mg/kg) | BPaUG8 |
| | | Bedaquiline (25 mg/kg)<br>Pretomanid (40 mg/kg)<br>Quabodepistat (14 mg/kg)<br>Sutezolid (100 mg/kg) | BPaQU |
| | | Bedaquiline (25 mg/kg)<br>Pretomanid (40 mg/kg)<br>Quabodepistat (14 mg/kg)<br>GSK-286 (35 mg/kg) | BPaQG2 |
| | | Bedaquiline (25 mg/kg)<br>Pretomanid (40 mg/kg)<br>Quabodepistat (14 mg/kg)<br>GSK-286 (35 mg/kg) | BQUG2 |
| | | Pretomanid (40 mg/kg)<br>Quabodepistat (14 mg/kg)<br>Sutezolid (100 mg/kg)<br>GSK-286 (35 mg/kg) | DQUG2 |
| | | Delamanid (2 mg/kg)<br>Quabodepistat (14 mg/kg)<br>Sutezolid (100 mg/kg)<br>GSK-286 (35 mg/kg) | PaQUG2 |
| E [iii,v] | Data submission to Critical Path Institute; TB-Platform for the Aggregation of Preclinical Experiments Data (TB-APEX) in progress<br><br>Data available at 10.5281/zenodo.21397999 | Bedaquiline (25 mg/kg)<br>Pretomanid (50 mg/kg)<br>Moxifloxacin (100 mg/kg)<br>Pyrazinamide (150 mg/kg) | BPaMZ |
| | | Bedaquiline (25 mg/kg)<br>Pretomanid (50 mg/kg) | BPa |
| | | Bedaquiline (25 mg/kg)<br>Pretomanid (50 mg/kg)<br>Sutezolid (50 mg/kg) | BPaU |
| | | Bedaquiline (25 mg/kg)<br>Pretomanid (50 mg/kg)<br>Linezolid (100 mg/kg) | BPaL |
| | | Bedaquiline (25 mg/kg)<br>Pretomanid (50 mg/kg)<br>TBD09 (200 mg/kg) | BPaT9 |
| | | Bedaquiline (25 mg/kg)<br>Pretomanid (50 mg/kg)<br>MK-3854 (200 mg/kg) | BPa3854 |
| | | Bedaquiline (25 mg/kg)<br>Pretomanid (50 mg/kg)<br>Moxifloxacin (100 mg/kg) | BPaM |
| | | Bedaquiline (25 mg/kg)<br>Pretomanid (50 mg/kg)<br>Moxifloxacin (100 mg/kg)<br>Sutezolid (50 mg/kg) | BPaMU |
| | | Bedaquiline (25 mg/kg)<br>Pretomanid (50 mg/kg)<br>TBD09 (200 mg/kg)<br>Moxifloxacin (100 mg/kg) | BPaT9M |
| F [iii,v] | | Bedaquiline (25 mg/kg) | BPaMZ |

| Dataset | Data source | Regimen | Abbr. |
|---|---|---|---|
| | Data submission to Critical Path Institute; TB-Platform for the Aggregation of Preclinical Experiments Data (TB-APEX) in progress<br><br>Data available at 10.5281/zenodo.21397999 | Pretomanid (50 mg/kg)<br>Moxifloxacin (100 mg/kg)<br>Pyrazinamide (150 mg/kg) | |
| | | Bedaquiline (25 mg/kg)<br>Pretomanid (50 mg/kg) | BPa |
| | | Bedaquiline (25 mg/kg)<br>Pretomanid (50 mg/kg)<br>Sutezolid (50 mg/kg) | BPaU |
| | | Bedaquiline (25 mg/kg)<br>Pretomanid (50 mg/kg)<br>Linezolid (100 mg/kg) | BPaL |
| | | Bedaquiline (25 mg/kg)<br>Pretomanid (50 mg/kg)<br>TBD11 (5 mg/kg) | BPaT1 |
| | | Bedaquiline (25 mg/kg)<br>Pretomanid (50 mg/kg)<br>GSK-286 (35 mg/kg) | BPaG2 |
| | | Bedaquiline (25 mg/kg)<br>Pretomanid (50 mg/kg)<br>Sutezolid (100 mg/kg)<br>GSK-286 (35 mg/kg) | BPaUG2 |
| | | Bedaquiline (25 mg/kg)<br>Pretomanid (50 mg/kg)<br>Sutezolid (50 mg/kg)<br>TBD11 (5 mg/kg) | BPaUT1 |
| G [iv,v] | Critical Path Institute; TB-Platform for the Aggregation of Preclinical Experiments Data (TB-APEX) Wave 2023, EV-TL-TBa23001 | Isoniazid (25 mg/kg)<br>Rifampin (10 mg/kg)<br>Pyrazinamide (150 mg/kg)<br>Ethambutol (100 mg/kg) | HRZE |
| | | Bedaquiline (25 mg/kg)<br>Pretomanid (100 mg/kg)<br>Moxifloxacin (100 mg/kg)<br>Pyrazinamide (150 mg/kg) | BPaMZ |
| | | Bedaquiline (25 mg/kg)<br>Delamanid (2 mg/kg)<br>Ganfeborole (8 mg/kg)<br>GSK-286 (35 mg/kg) | BDG6G2 |
| | | Bedaquiline (25 mg/kg)<br>Quabodepistat (14 mg/kg)<br>Ganfeborole (8 mg/kg)<br>GSK-286 (35 mg/kg) | BQG6G2 |
| | | Bedaquiline (25 mg/kg)<br>Quabodepistat (14 mg/kg)<br>Sutezolid (100 mg/kg)<br>Ganfeborole (8 mg/kg) | BQUG6 |
| | | Bedaquiline (25 mg/kg)<br>Pretomanid (40 mg/kg)<br>Ganfeborole (8 mg/kg)<br>GSK-286 (35 mg/kg) | BPaG6G2 |
| | | Bedaquiline (25 mg/kg)<br>Pretomanid (40 mg/kg)<br>Quabodepistat (14 mg/kg)<br>Ganfeborole (8 mg/kg) | BPaQG6 |
| | | Bedaquiline (25 mg/kg)<br>Sutezolid (100 mg/kg)<br>Ganfeborole (8 mg/kg)<br>GSK-286 (35 mg/kg) | BUG6G2 |
| | | Delamanid (2 mg/kg)<br>Quabodepistat (14 mg/kg)<br>Ganfeborole (8 mg/kg)<br>GSK-286 (35 mg/kg) | DQG6G2 |
| | | Delamanid (2 mg/kg)<br>Quabodepistat (14 mg/kg)<br>Sutezolid (100 mg/kg)<br>Ganfeborole (8 mg/kg) | DQUG6 |
| | | Delamanid (2 mg/kg)<br>Sutezolid (100 mg/kg)<br>Ganfeborole (8 mg/kg) | DUG6G2 |

| Dataset | Data source | Regimen | Abbr. |
|---|---|---|---|
| | | GSK-286 (35 mg/kg) | |
| | | Quabodepistat (14 mg/kg)<br>Sutezolid (100 mg/kg)<br>Ganfeborole (8 mg/kg)<br>GSK-286 (35 mg/kg) | QUG6G2 |
| | | Pretomanid (40 mg/kg)<br>Quabodepistat (14 mg/kg)<br>Ganfeborole (8 mg/kg)<br>GSK-286 (35 mg/kg) | PaQG6G2 |
| | | Pretomanid (40 mg/kg)<br>Quabodepistat (14 mg/kg)<br>Sutezolid (100 mg/kg)<br>Ganfeborole (8 mg/kg) | PaQUG6 |
| | | Pretomanid (40 mg/kg)<br>Sutezolid (100 mg/kg)<br>Ganfeborole (8 mg/kg)<br>GSK-286 (35 mg/kg) | PaUG6G2 |
| H [iv,v] | Critical Path Institute;<br>TB-Platform for the Aggregation of Preclinical Experiments Data (TB-APEX) RMM2023_2, EV-TL-TBa23007 | Bedaquiline (25 mg/kg)<br>Pretomanid (100 mg/kg)<br>Moxifloxacin (100 mg/kg)<br>Pyrazinamide (150 mg/kg) | BPaMZ |
| | | Isoniazid (10 mg/kg)<br>Rifapentine (10 mg/kg)<br>Moxifloxacin (100 mg/kg)<br>Pyrazinamide (150 mg/kg) | HPZM |
| | | Sorfequiline (TBAJ-876) (6.25 mg/kg)<br>Delamanid (2 mg/kg)<br>Sutezolid (100 mg/kg)<br>Ganfeborole (8 mg/kg) | S8DUG6 |
| | | Sorfequiline (TBAJ-876) (6.25 mg/kg)<br>Pretomanid (40 mg/kg)<br>Sutezolid (100 mg/kg)<br>Ganfeborole (8 mg/kg) | S8PaUG6 |
| | | Bedaquiline (25 mg/kg)<br>Pretomanid (40 mg/kg)<br>Sutezolid (100 mg/kg)<br>Ganfeborole (8 mg/kg) | BPaUG6 |
| | | Bedaquiline (25 mg/kg)<br>Pretomanid (40 mg/kg)<br>Sutezolid (100 mg/kg)<br>GSK-830 (5 mg/kg) | BPaUG8 |
| I [vi] | Critical Path Institute;<br>TB-Platform for the Aggregation of Preclinical Experiments Data (TB-APEX) RMM2024_1, EV-TL-TBa24001 | Bedaquiline (25 mg/kg)<br>Pretomanid (100 mg/kg)<br>Moxifloxacin (100 mg/kg)<br>Pyrazinamide (150 mg/kg) | BPaMZ |
| | | Isoniazid (10 mg/kg)<br>Rifapentine (10 mg/kg)<br>Moxifloxacin (100 mg/kg)<br>Pyrazinamide (150 mg/kg) | HPZM |
| | | Sorfequiline (TBAJ-876) (6.25 mg/kg)<br>Pretomanid (40 mg/kg)<br>TBD11 (5 mg/kg) | S8PaT1 |
| | | Sorfequiline (TBAJ-876) (6.25 mg/kg)<br>Pretomanid (40 mg/kg)<br>Quabodepistat (14 mg/kg)<br>TBD11 (5 mg/kg) | S8PaQT1 |
| | | Sorfequiline (TBAJ-876) (6.25 mg/kg)<br>Pretomanid (40 mg/kg)<br>TBD09 (200 mg/kg) | S8PaT9 |
| | | Sorfequiline (TBAJ-876) (6.25 mg/kg)<br>Quabodepistat (14 mg/kg)<br>TBD09 (200 mg/kg) | S8QT9 |
| | | Sorfequiline (TBAJ-876) (6.25 mg/kg)<br>Pretomanid (40 mg/kg)<br>Quabodepistat (14 mg/kg)<br>TBD09 (200 mg/kg) | S8PaQT9 |
| | | Sorfequiline (TBAJ-876) (6.25 mg/kg)<br>Quabodepistat (14 mg/kg) | S8QT1T9 |

| Dataset | Data source | Regimen | Abbr. |
|---|---|---|---|
| | | TBD11 (5 mg/kg)<br>TBD09 (200 mg/kg) | |
| | | Sorfequiline (TBAJ-876) (6.25 mg/kg)<br>Pretomanid (40 mg/kg)<br>Sutezolid (100 mg/kg) | S8PaU |
| | | Sorfequiline (TBAJ-876) (6.25 mg/kg)<br>Pretomanid (40 mg/kg)<br>GSK-286 (35 mg/kg) | S8PaG2 |
| | | Sorfequiline (TBAJ-876) (6.25 mg/kg)<br>Pretomanid (40 mg/kg)<br>Sutezolid (100 mg/kg)<br>GSK-286 (35 mg/kg) | S8PaUG2 |
| | | Sorfequiline (TBAJ-876) (6.25 mg/kg)<br>Pretomanid (40 mg/kg)<br>Quabodepistat (14 mg/kg) | S8PaQ |
| | | Sorfequiline (TBAJ-876) (6.25 mg/kg)<br>Pretomanid (40 mg/kg)<br>Quabodepistat (14 mg/kg)<br>GSK-286 (35 mg/kg) | S8PaQG2 |
| | | Sorfequiline (TBAJ-876) (6.25 mg/kg)<br>Quabodepistat (14 mg/kg)<br>GSK-286 (35 mg/kg)<br>TBD09 (200 mg/kg) | S8QG2T9 |

**Colorado State University (A,B)**: Bedaquiline (LKT Laboratories, St. Paul, MN, USA); Delamanid (Otsuka Pharmaceutical Co., Ltd., Chiyoda-ku, Tokyo, Japan); Ethambutol (Millipore-Sigma, Burlington, MA, USA); Isoniazid (Millipore-Sigma, Burlington, MA, USA); Linezolid (LKT Laboratories, St. Paul, MN, USA); Pretomanid (ChemShuttle, Burlingame, CA, USA); Pretomanid (ChemShuttle, Burlingame, CA, USA); Pyrazinamide (Acros Organics, Geel, Belgium); Quabodepistat (Otsuka Pharmaceutical Co., Ltd., Chiyoda-ku, Tokyo, Japan); Rifampin (Millipore-Sigma, Burlington, MA, USA); Sutezolid (RTI International, Research Triangle Park, NC, USA) **Evotec (D, G, H, I)**: Bedaquiline (Johnson & Johnson Innovative Medicine, Titusville, NJ, USA); Delamanid (Otsuka Pharmaceutical Co., Ltd., Chiyoda-ku, Tokyo, Japan); Ethambutol (Millipore-Sigma, Burlington, MA, USA); Ganfeborole (GSK, London, United Kingdom); GSK-286 (GSK, London, United Kingdom); GSK-830 (GSK, London, United Kingdom); Isoniazid (Millipore-Sigma, Burlington, MA, USA); Isoniazid (Millipore-Sigma, Burlington, MA, USA); Moxifloxacin (LKT Laboratories, St. Paul, MN, USA); Pretomanid (TB Alliance, New York, NY, USA); Pyrazinamide (Millipore-Sigma, Burlington, MA, USA); Quabodepistat (Otsuka Pharmaceutical Co., Ltd., Chiyoda-ku, Tokyo, Japan); Rifampin (Millipore-Sigma, Burlington, MA, USA); Rifapentine (Millipore-Sigma, Burlington, MA, USA); Sorfequiline (TBAJ-876) (TB Alliance, New York, NY, USA); Sutezolid (TB Alliance, New York, NY, USA); TBD09 (Gates MRI, Cambridge, MA, USA); TBD11 (Gates MRI, Cambridge, MA, USA). **Johns Hopkins University (C, E, F):** Bedaquiline (Biosynth, Staad, Switzerland); Ethambutol (Millipore-Sigma, Burlington, MA, USA); GSK-286 (GSK, London, United Kingdom); Isoniazid (Millipore-Sigma, Burlington, MA, USA); Linezolid (Millipore-Sigma, Burlington, MA, USA); MK-3854 (Merck (via GMRI), Rahway, NJ, USA); Moxifloxacin (Biosynth, Staad, Switzerland); Pretomanid (TB Alliance, New York, NY, USA); Pyrazinamide (Acros Organics, Geel, Belgium); Rifabutin (Biosynth, Staad, Switzerland); Rifampin (Millipore-Sigma, Burlington, MA, USA); Rifapentine (Sanofi-Aventis, Paris, France); Sutezolid (TB Alliance, New York, NY, USA); TBD09 (Merck (via GMRI), Rahway, NJ, USA); TBD11 (Calibr (via GMRI), La Jolla, CA, USA)

**Table S2.** Univariable iteration 2 model development results with predictors listed with their summary statistics for the relapse-free and relapsing mice and with their model statistics including odds ratio (OR) and p-value. Italics: Odds ratio (OR) rounded to 1 when rounding to 7 significant digits (but coefficient not equal to zero). CI – Confidence Interval; sd – standard deviation.

| **Predictor** | **No relapse (mean (sd))** | **Relapse (mean (sd))** | **OR** | **OR (95% CI)** | **P-value** | **missing** |
|---|---|---|---|---|---|---|
| Treatment duration | 79.13 (26.74) | 56.48 (24.82) | 0.967 | (0.962-0.972) | 4.15e-52 | 0 |
| log10 RS ratio change at day 28 | 1.79 (0.45) | 1.51 (0.63) | 0.370 | (0.295-0.463) | 1.00e-20 | 0 |
| log10 CFU/lung change at day 28 | 4.85 (1.68) | 4.07 (1.63) | 0.755 | (0.705-0.808) | 6.44e-17 | 0 |
| log10 RS ratio change at day 14 | 1.55 (0.47) | 1.28 (0.62) | 0.411 | (0.332-0.51) | 1.08e-17 | 0 |
| Washout | 85.35 (3.09) | 84.13 (3.12) | 0.871 | (0.836-0.908) | 6.01e-13 | 0 |
| log10 CFU/lung change at day 14 | 2.11 (0.86) | 1.82 (0.79) | 0.658 | (0.576-0.752) | 4.32e-10 | 0 |
| Intranasal route (relative to aerosol) | - | - | 1.983 | (1.542-2.552) | 6.66e-08 | 0 |
| RS ratio delta between day 14 and 28 | 5.65 (11.32) | 14.58 (31.99) | 1.025 | (1.015-1.034) | 1.64e-13 | 0 |
| RS ratio at day 14 | 14.08 (37.74) | 57.82 (160.68) | 1.010 | (1.006-1.015) | 3.24e-17 | 0 |
| RS ratio at day 28 | 8.43 (32.2) | 43.24 (147.44) | 1.019 | (1.011-1.027) | 1.51e-15 | 0 |
| RS ratio change at day 14 | 256.73 (44.31) | 218.1 (161.3) | 0.995 | (0.993-0.997) | 5.64e-11 | 0 |
| CFU/lung at day 28 | 56462.75 (196585.02) | 130264.87 (359559.58) | 1.000001 | (1.0000005-1.0000015) | 2.20e-06 | 0 |
| log10 RS ratio baseline | 2.43 (0.04) | 2.44 (0.03) | 498.761 | (26.141-9516.353) | 3.25e-05 | 0 |
| CFU/lung change at day 14 | *41307235.83 (21129770.47)* | *36657898.3 (19404230.75)* | *1.000* | *(1-1)* | *3.44e-05* | *0* |
| RS ratio baseline | 270.81 (24.23) | 275.92 (21.79) | 1.010 | (1.005-1.014) | 5.91e-05 | 0 |
| Erdman (relative to H37Rv) strain | - | - | 0.612 | (0.481-0.779) | 6.02e-05 | 0 |
| RS ratio change at day 28 | 262.38 (40.25) | 232.68 (148.26) | 0.996 | (0.994-0.998) | 7.56e-08 | 0 |
| CFU/lung change at day 28 | *42887322.46 (22193442.32)* | *38811237.87 (20948634.7)* | *1.000* | *(1-1)* | *6.32e-04* | *0* |
| CFU/lung baseline | *42943785.21 (22240794.4)* | *38941502.74 (21095127.68)* | *1.000* | *(1-1)* | *8.35e-04* | *0* |
| CFU/lung at day 14 | 1636549.38 (3064335.95) | 2283604.44 (3915645.91) | 1.0000001 | (1.0000000-1.0000001) | 8.17e-04 | 0 |
| log10 CFU/lung baseline | 7.57 (0.24) | 7.53 (0.23) | 0.497 | (0.315-0.784) | 2.59e-03 | 0 |
| Oxazolidinone present in regimen | - | - | 0.888 | (0.715-1.102) | 2.80e-01 | 0 |

**Table S3.** Univariable iteration 3 model development results with predictors listed with their summary statistics for the relapse-free and relapsing mice and with their model statistics including odds ratio (OR) and p-value. Italics: Odds ratio (OR) rounded to 1 when rounding to 7 significant digits (but coefficient not equal to zero). CI – Confidence Interval; sd – standard deviation.

| **Predictor** | **No relapse (mean (sd))** | **Relapse (mean (sd))** | **OR** | **OR (95% CI)** | **P-value** | **missing** |
|---|---|---|---|---|---|---|
| Treatment duration | 76.35 (25.78) | 54.97 (23.63) | 0.966 | (0.962-0.97) | 7.33e-70 | 0 |
| log10 RS ratio change at day 28 | 1.76 (0.46) | 1.24 (0.83) | 0.29 | (0.243-0.345) | 5.62e-59 | 0 |
| log10 CFU/lung change at day 28 | 5.23 (1.86) | 4.33 (1.65) | 0.748 | (0.708-0.79) | 2.83e-27 | 0 |
| log10 RS ratio change at day 14 | 1.53 (0.47) | 1.01 (0.83) | 0.301 | (0.253-0.358) | 9.40e-57 | 0 |
| Washout | 85.06 (2.79) | 84.09 (2.52) | 0.855 | (0.819-0.892) | 1.05e-15 | 0 |
| log10 CFU/lung change at day 14 | 2.37 (0.96) | 2.14 (0.96) | 0.782 | (0.71-0.861) | 4.62e-07 | 25 |
| Intranasal route (relative to aerosol) | - | - | 2.154 | (1.785-2.598) | 5.87e-16 | 0 |
| RS ratio delta between day 14 and 28 | 6.4 (20.26) | 56.86 (116.39) | 1.02 | (1.015-1.025) | 1.86e-50 | 0 |
| RS ratio at day 14 | 15.49 (43.03) | 171.31 (369.64) | 1.011 | (1.008-1.013) | 4.14e-59 | 0 |
| RS ratio at day 28 | 9.09 (30.27) | 114.46 (268.91) | 1.023 | (1.018-1.029) | 7.95e-58 | 0 |
| RS ratio change at day 14 | 249.88 (49.11) | 95.65 (374.69) | 0.993 | (0.992-0.995) | 5.73e-50 | 0 |
| CFU/lung at day 28 | *44465.9 (175490.83)* | *98032.11 (302829.49)* | *1* | *(1-1)* | *2.37e-06* | *0* |
| log10 RS ratio baseline | 2.42 (0.04) | 2.43 (0.03) | 9.098 | (0.725-114.13) | 8.66e-02 | 0 |
| CFU/lung change at day 14 | 39952155.38 (18942635.64) | 37024277.1 (15976261.75) | 1 | (1-1) | 3.61e-04 | 25 |
| RS ratio baseline | 265.37 (23.91) | 266.96 (21.64) | 1.003 | (0.999-1.007) | 1.34e-01 | 0 |
| Erdman (relative to H37Rv) strain | | | 0.513 | (0.408-0.645) | 8.81e-09 | 0 |
| RS ratio change at day 28 | 256.28 (38.68) | 152.5 (273.5) | 0.992 | (0.989-0.994) | 2.11e-40 | 0 |
| CFU/lung change at day 28 | *41203403 (19965583.46)* | *38595136.13 (17023600.46)* | *1* | *(1-1)* | *2.52e-03* | *0* |
| CFU/lung baseline | *41247868.9 (20010532.17)* | *38693168.24 (17143201.53)* | *1* | *(1-1)* | *3.22e-03* | *0* |
| CFU/lung at day 14 | *1295713.52 (2789355.64)* | *1652303.29 (3356582.84)* | *1* | *(1-1)* | *1.41e-02* | *25* |
| log10 CFU/lung baseline | 7.56 (0.22) | 7.55 (0.19) | 0.676 | (0.429-1.066) | 9.19e-02 | 0 |
| Oxazolidinone present in regimen | - | - | 0.919 | (0.765-1.104) | 3.67e-01 | 0 |

**Table S4**. Multivariable iteration 2 model development steps with the included parameter or deleted parameter and p-value per step for the forward inclusion or backward deletion, respectively. Oxazolidinone present in regimen was included a priori for predictive purposes.

| **Step** | **Parameter** | **p-value** |
|---|---|---|
| Forward inclusion | | |
| 1 | *Oxazolidinone present in regimen* | *A priori* |
| 2 | Treatment duration | 9.15 e-54 |
| 3 | RS log10 change at day 28 | 3.81 e-95 |
| 4 | CFU log10 change at day 28 | 7.64 e-39 |
| 5 | Strain | 1.32 e-05 |
| 6 | CFU log10 change at day 14 | 9.61 e-06 |
| 7 | Infection route | 1.64 e-08 |
| 8 | RS baseline | 1.76 e-05 |
| 9 | CFU log10 baseline | 3.98 e-03 |
| 10 | RS log10 change at day 14 | 4.40 e-03 |
| Backward deletion | | |
| 11 | RS log10 change at day 28 | 0.2449 |

**Table S5**. Multivariable iteration 3 model development steps with the included parameter or deleted parameter and p-value per step for the forward inclusion or backward deletion, respectively. Oxazolidinone present in regimen was included a priori for predictive purposes.

| Step | Parameter | p-value |
|---|---|---|
| Forward inclusion | | |
| 1 | *Oxazolidinone present in regimen* | *A priori* |
| 2 | Treatment duration | 3.93e-71 |
| 3 | RS log10 change at day 14 | 6.03e-153 |
| 4 | CFU log10 change at day 28 | 4.73e-50 |
| 5 | Strain | 0.00023 |
| 6 | CFU log10 change at day 14 | 5.26e-06 |
| 7 | Infection route | 6.36e-06 |
| 8 | CFU log10 baseline | 0.0011 |
| 9 | RS baseline | 0.00012 |
| | Backward deletion | |
| 11 | *All backward deletion p<0.0005* | |

**Figure S1.** Forest plot of the final model. Minimal changes in the coefficient estimates are shown in this final model structure for the training datasets 2 (yellow), training dataset 3 (burgundy), and once re-estimated for the external validation dataset 3 (teal). CI = confidence interval, CFU = colony forming units, OR = odds ratio

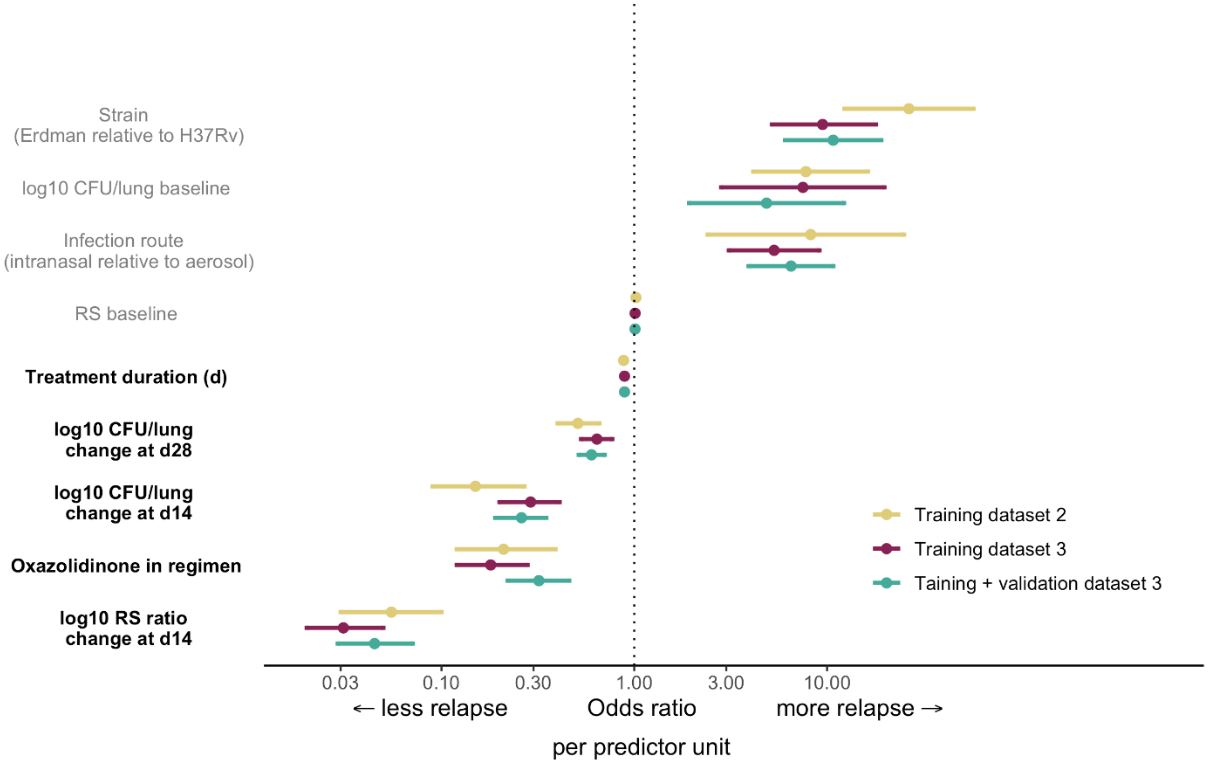

**Figure S2.** Receiver Operator Curve (ROC) for each of the multivariable step models (Table S4-5) of model iteration 2 (**A**) and 3 (**B**) and their area under the ROC (AUROC). The final model of iteration 2 with AUROC = 0.929 is the model after backward deletion as listed in step 11 of Table S3.

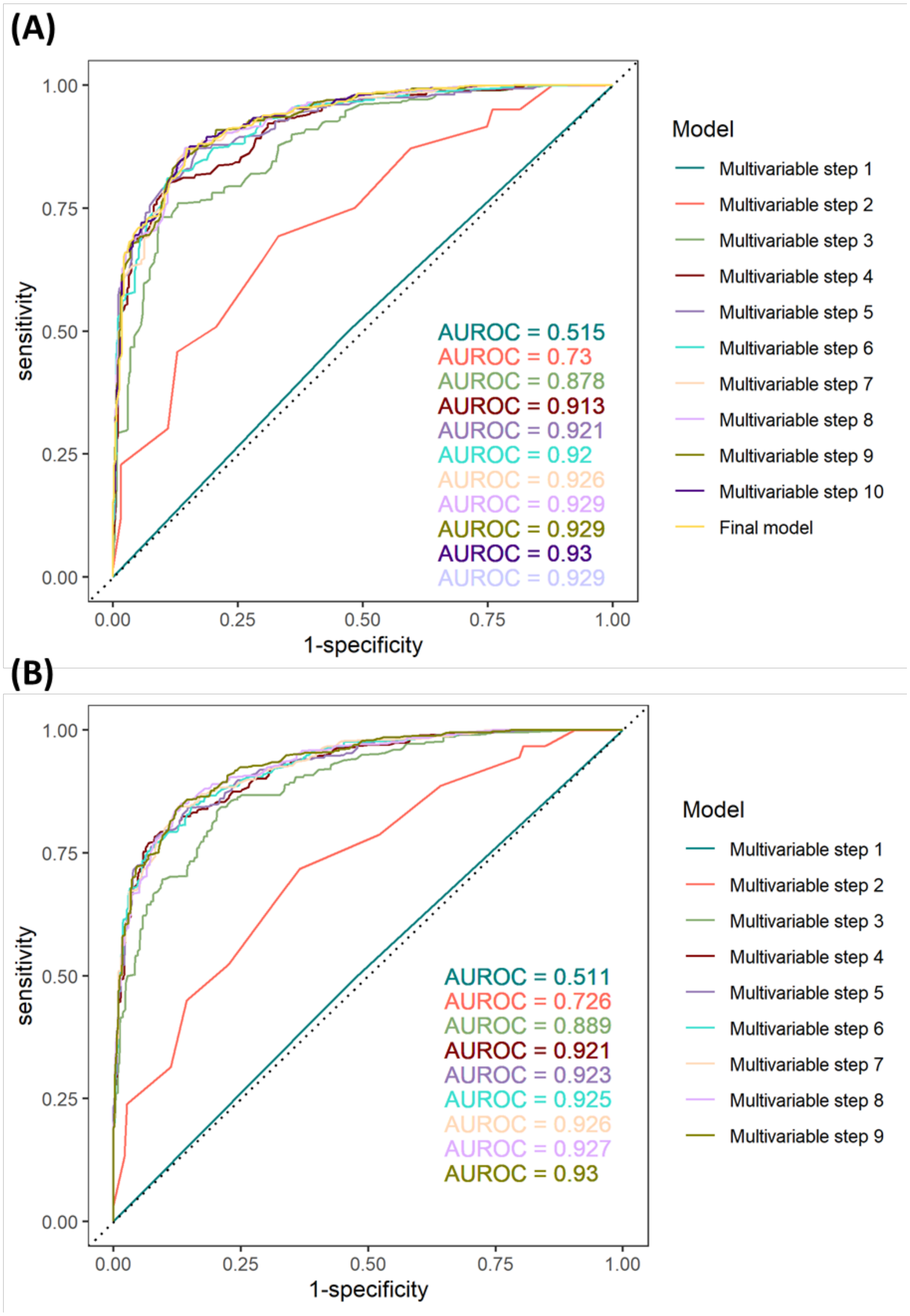

**Figure S3.** Model predicted relapse over time of training dataset 2 (**A**) and validation dataset 2 (**B**) by the iteration 2 model. Observed proportions of relapsing mice are shown in symbols, lines and intervals show median and 95% prediction interval, colors show the experiment. Regimens are defined in **Table 1.**

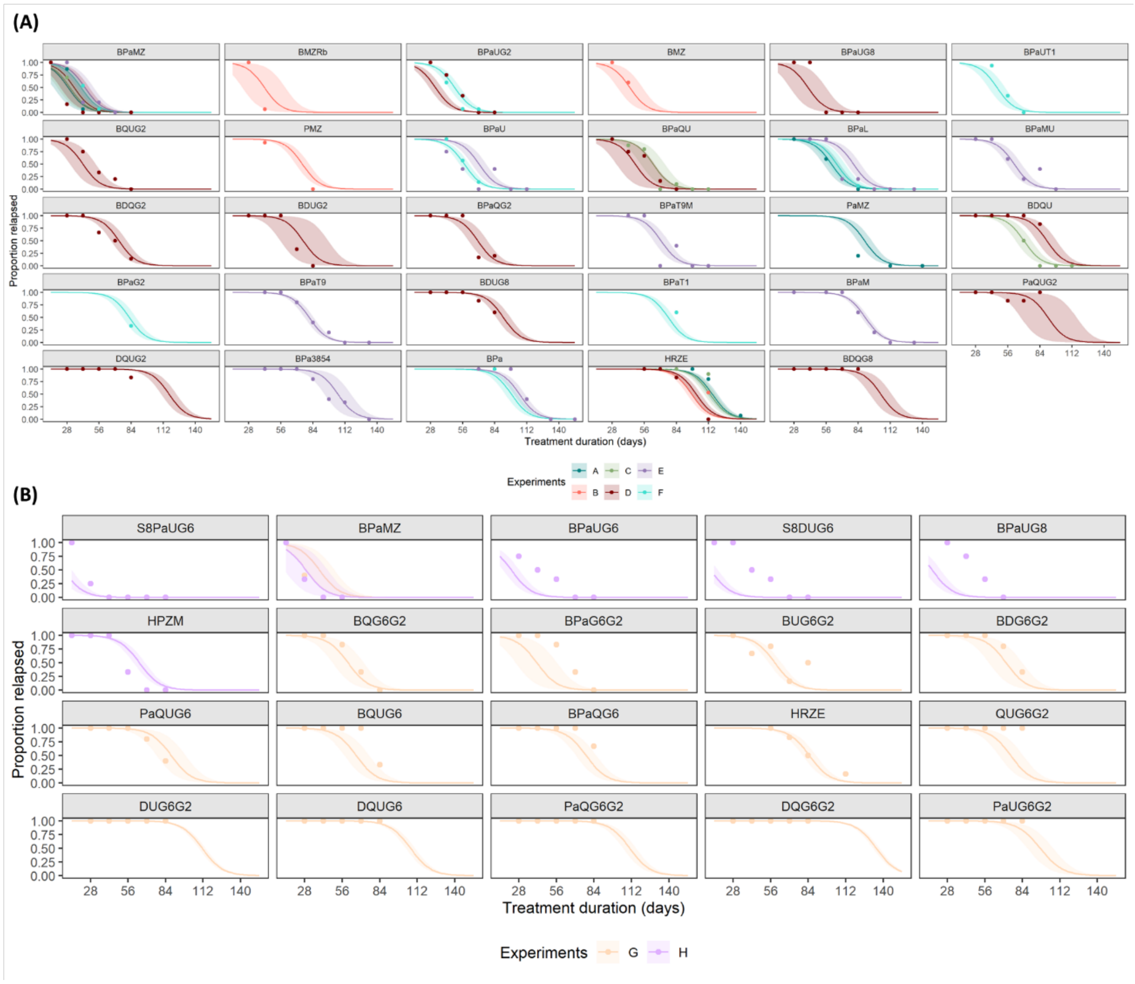

**Figure S4.** Forest plot of predicted timepoint at which 95% of mice are relapse-free (T95) based on the final model (iteration 3) with baseline correction and for H37Rv strain infection through the aerosol route for the training (**A**) and validation (**B**) datasets. CI = confidence interval, regimens are defined in Table 1.

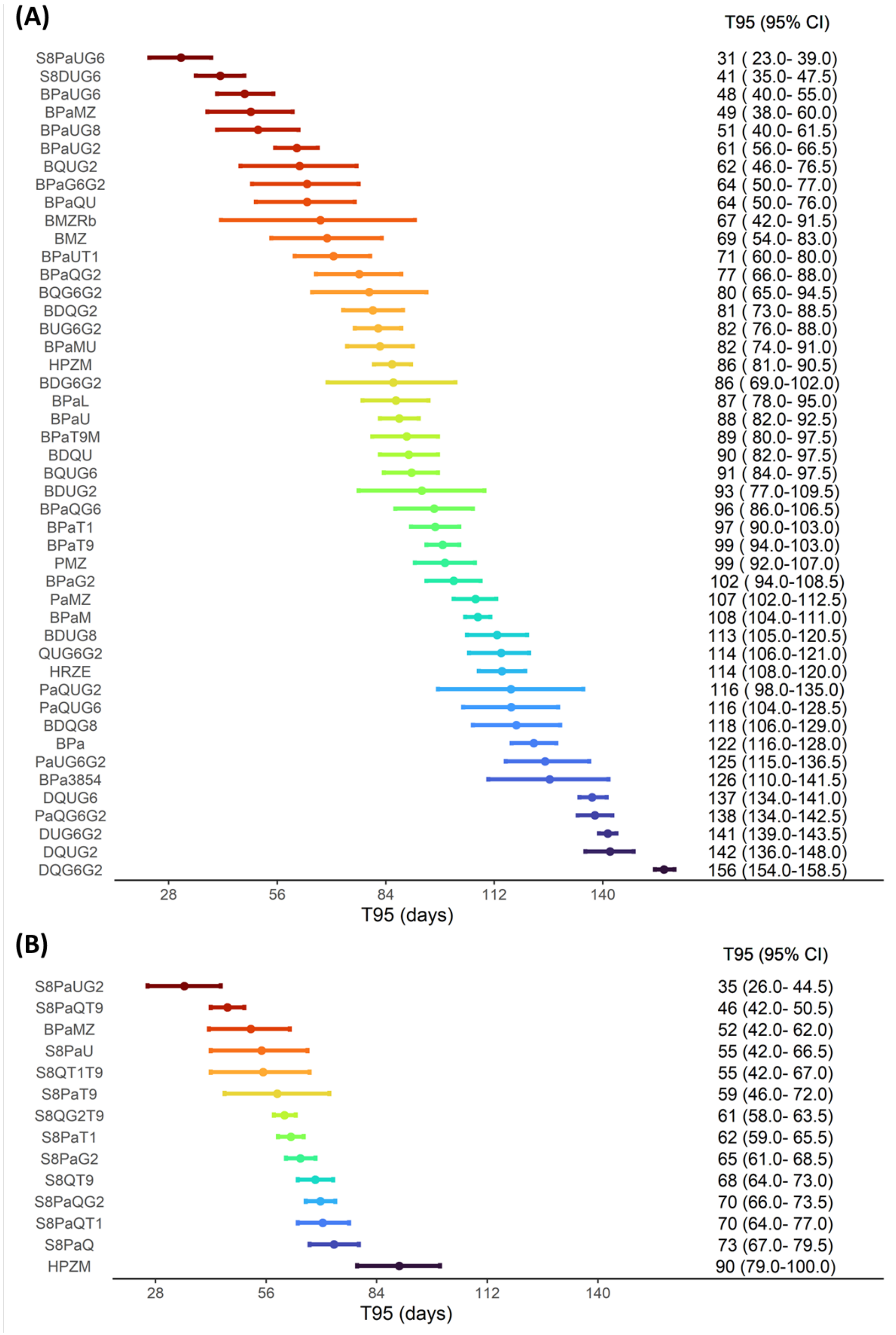

**Figure S5.** Forest plot of predicted timepoint at which 95% of mice are relapse-free (T95) based on the final model (iteration 3) with baseline correction and for Erdman strain infection through the aerosol route for the training (**A**) and validation (**B**) datasets. CI = confidence interval, regimens are defined in Table 1.

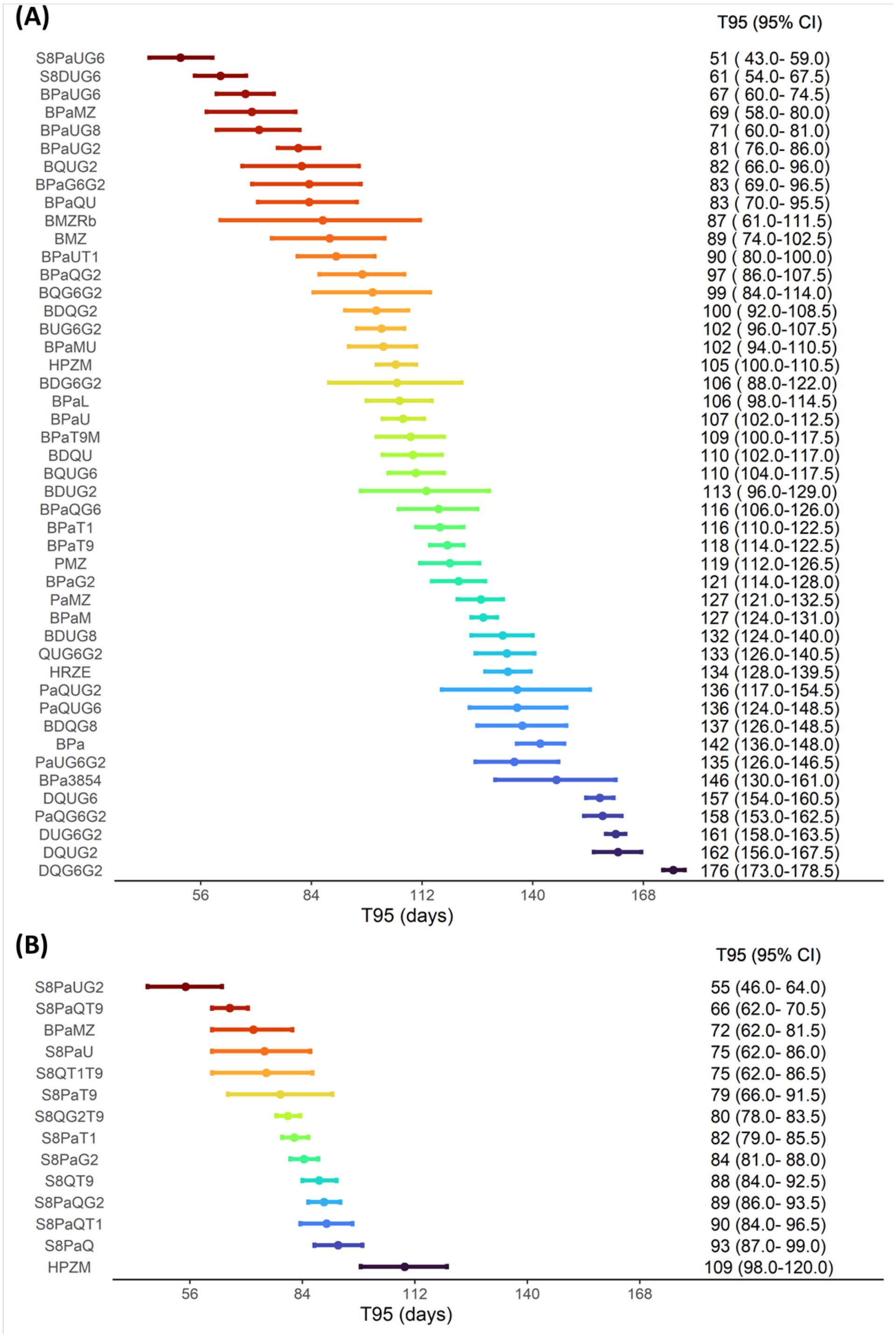

**Figure S6**. Forest plot of predicted timepoint at which 95% of mice are relapse-free (T95) based on the final model (iteration 3) with baseline correction and for H37Rv strain infection through the intranasal route for the training (**A**) and validation (**B**) datasets. CI = confidence interval, regimens are defined in **Table 1.**

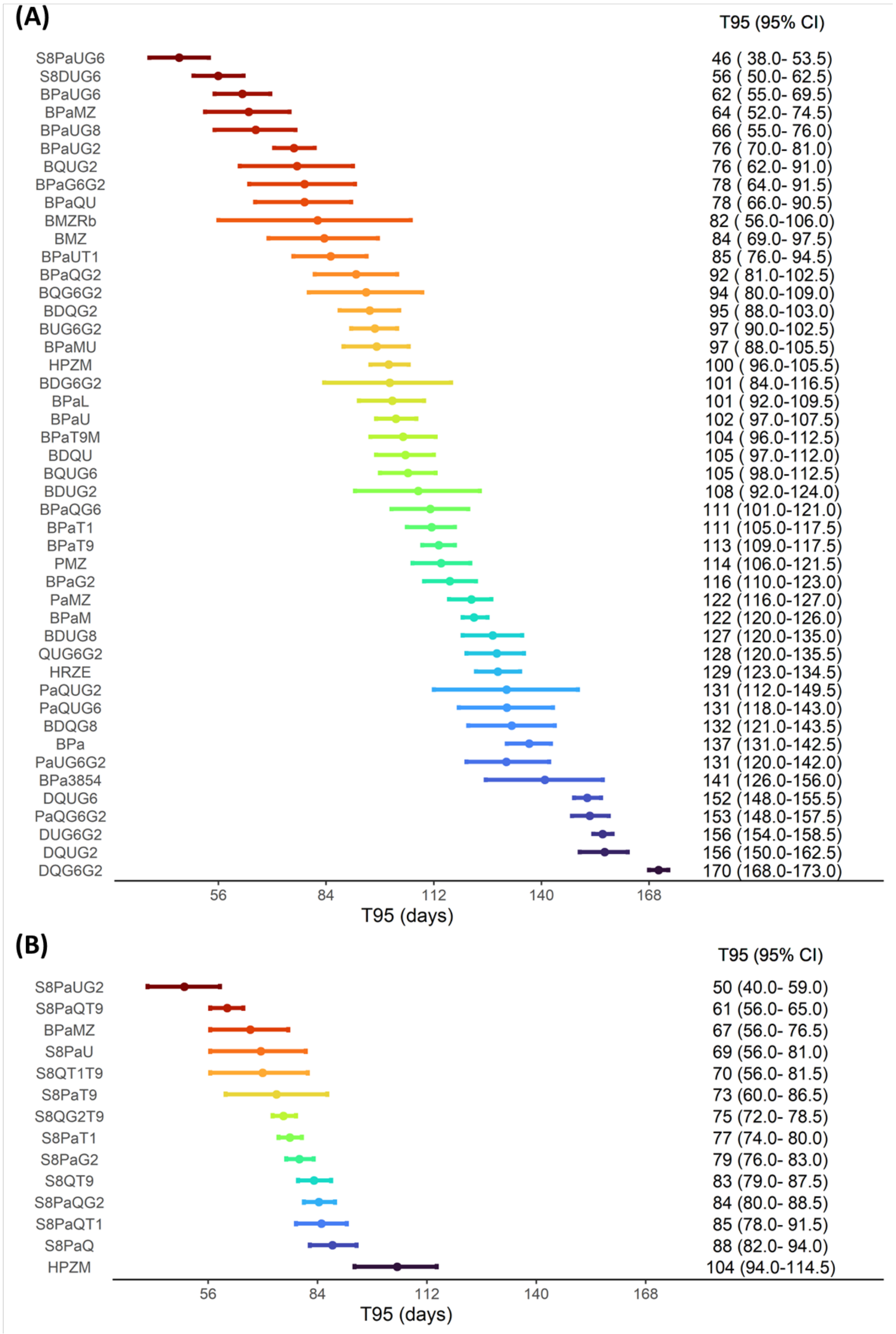

**Figure S7.** Model predicted T95 (rainbow colors) for the final model (iteration 3) as function of the CFU and RS ratio changes at 14 and 28 days, with 2, 3, and 4 month treatment duration shown as dotted, solid, and dashed diagonal lines, respectively. Left panel shows the changes in RS ratio at day 14 and the changes in CFU at day 28 with changes in CFU at day 14 constant at the median, middle panel shows the changes in RS ratio at day 14 and the changes in CFU at day 14 with changes in CFU at day 28 constant at the median, right panel shows the changes in CFU at day 14 and the changes in CFU at day 28 with changes in RS ratio at day 14 constant at the median without (**A**) and with (**B**) the observed values as symbols for all regimens in the training (circles) and validation (triangle) datasets. Prediction is baseline-corrected and shown for H37Rv infection by aerosol route.

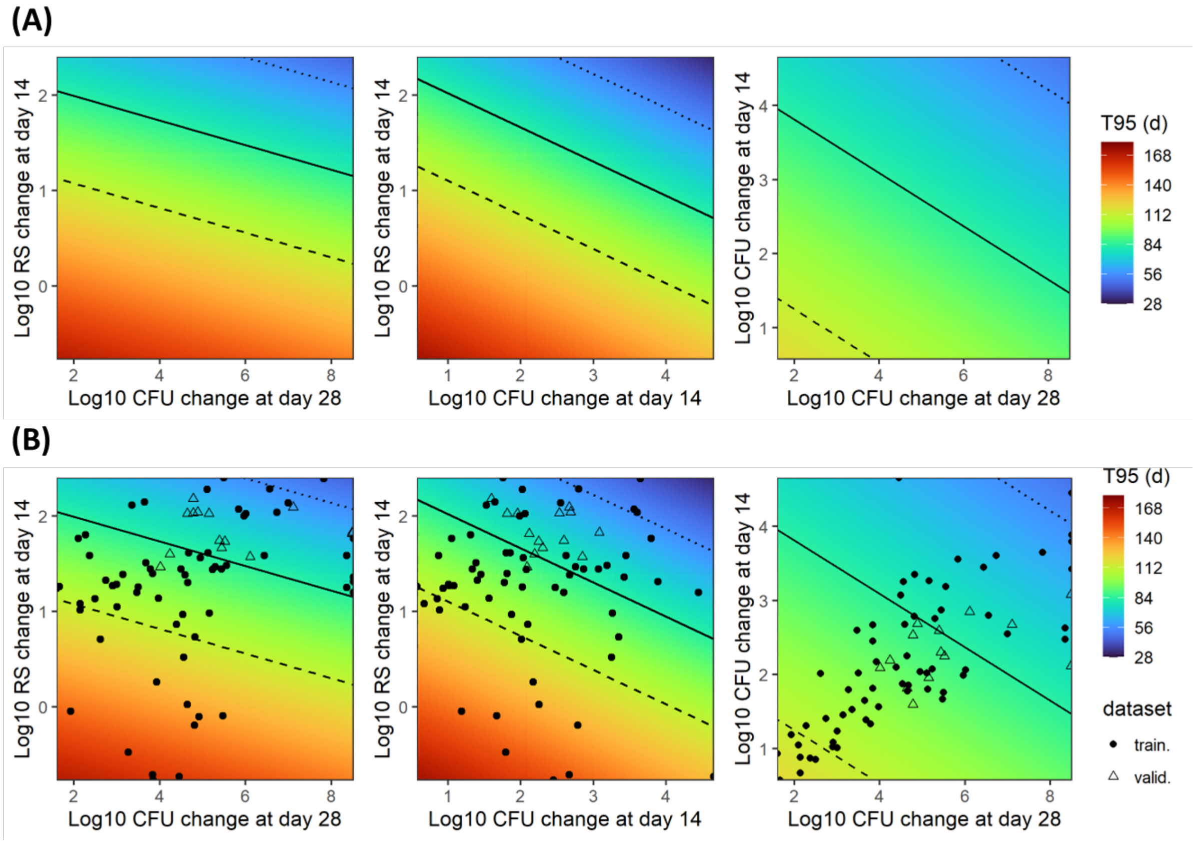
